# Human Pumilio proteins use fuzzy multivalent hydrophobic interactions to recruit the CCR4–NOT deadenylase complex to repress mRNAs

**DOI:** 10.64898/2025.12.18.695197

**Authors:** Elise B. Dunshee, Brenna A. Saladin, David J. Turner, Chen Qiu, Robert C. Dutcher, Jason G. Williams, Joshua Corbo, Olivia R. Wolcott, Amanda J. Korte, Rebecca J. Haugen, Traci M. Tanaka Hall, Eugene Valkov, Aaron C. Goldstrohm

**Author notes:** For correspondence: Aaron Goldstrohm.

## Abstract

Pumilio (PUM) proteins are conserved RNA-binding proteins that control mRNAs involved in development, proliferation, and stem cell differentiation. Human PUM1 and PUM2 repress targets by recruiting the CCR4–NOT deadenylase complex through a metazoan-specific N-terminal repression domain (RD3), which is predicted to be intrinsically disordered. Here we dissect RD3 using cell-based reporter assays, protein interaction assays with recombinant proteins, and crosslinking mass spectrometry. We identify multiple short RD3 peptides that are sufficient for repression and bind directly to the C-terminal NOT module of CCR4–NOT, comprising CNOT1, CNOT2, and CNOT3 subunits. Crosslinking reveals numerous mutually exclusive contacts between RD3 and the NOT module, consistent with a multivalent “fuzzy” binding mode in which interactions are not defined by a single sequence or structure. Sequence scrambling shows that the linear amino acid order of RD3 is dispensable, whereas its physicochemical composition, in particular distributed aliphatic and aromatic residues, is essential for repression and CCR4–NOT binding. These findings support a model in which low-affinity, multivalent interactions between intrinsically disordered regions (IDRs) and effector complexes, governed by amino acid composition rather than precise sequence, underlie robust PUM-mediated repression, and exemplify general principles by which IDRs recruit the CCR4–NOT complex to regulate gene expression.

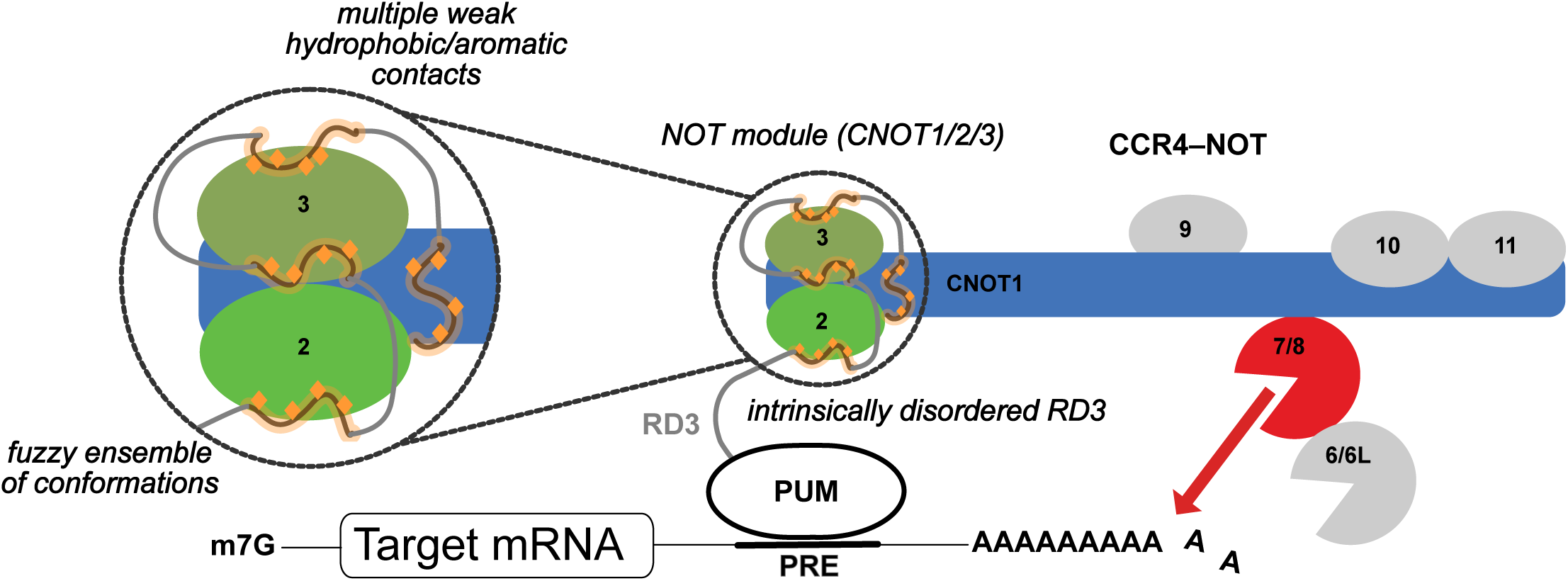
Graphical Abstract.

## Introduction

Regulation of gene expression is critical for orchestrating complex biological processes such as embryogenesis and cellular differentiation. Post-transcriptional control by RNA-binding proteins (RBPs) is a central node in this regulation, as RBPs govern mRNA fate by influencing transcript abundance and processing.^1–4^ Pumilio (PUM) proteins are model RBPs and essential regulators throughout eukaryotes.^5,6^ Humans express two PUM paralogs, PUM1 and PUM2, that bind thousands of RNAs in a sequence-specific manner with overlapping RNA-binding specificity and regulatory functions.^6–12^ Their targets include transcripts involved in neurodevelopment, cell-cycle control, proliferation, stem cell differentiation, and signaling pathways,^11^ and PUM dysfunction has been implicated in the progression of cancer^13–20^ and neurological disease.^21–24^

The PUM family is defined by a conserved C-terminal RNA-binding domain (RBD) that binds to specific mRNAs by recognizing an eight-nucleotide Pumilio response element (PRE; 5′ UGUANAUA) with high affinity and specificity.^10,12,25–27^ Hundreds of direct target mRNAs that are bound and regulated by PUM1 and PUM2 have been identified.^8–12^ Mechanistically, PUMs repress gene expression by accelerating decay of target mRNAs.^7,11,12,28^

PUMs from insects to humans contain large, unique N-terminal extensions. Functional analysis showed that these regions confer the major repressive activity of *Drosophila* Pumilio and human PUM1 and PUM2.^28,29^ Based on sequence conservation and functional analysis, the N-terminal regions were subdivided into three repression domains (RD1-3) interspersed by two Pumilio conserved motifs (PCMa and PCMb).^6,29–31^ In contrast to the structured RBD, the N-terminal regions, particularly the RDs, are poorly conserved and lack predicted structure.^6^ While all three RDs contribute to repression in *Drosophila* Pumilio,^29,30^ functional analysis of PUM1 and PUM2 indicated that the third repression domain, RD3, provides the dominant inhibitory activity.^28^

The PUM N-terminal RDs promote mRNA destruction by directly recruiting the CCR4–NOT deadenylase complex, which shortens poly(A) tails, triggering translational repression and mRNA decay.^7,28,30,31^ Thus, PUMs exhibit a modular architecture in which the RBD binds target mRNAs and the N-terminal RDs recruit CCR4–NOT to accelerate destruction of the mRNAs. In *Drosophila*, each Pumilio RD binds the eight-subunit CCR4–NOT complex.^30^ The points of contact of RD3 were narrowed to three subunits of the deadenylase complex (CNOT1, CNOT2 and CNOT3).^31^ Conserved motifs that confer repressive activity and CCR4–NOT binding were mapped within *Drosophila* RD3 and interpreted as short linear interaction motifs (SLIMs).^31^ For human PUM1 and PUM2, biochemical analysis identified direct binding of RD3 to the NOT module, which contains the C-terminal region of CNOT1 and the CNOT2 and CNOT3 subunits,^28^ but the molecular basis of this interaction remained unknown.

RD3 from both human PUMs has the hallmarks of an intrinsically-disordered region (IDR). Overall, the N-terminal regions lack homology to known structural domains and exhibit rapid evolutionary changes.^6^ Moreover, sequence analysis of RD3 did not identify predicted secondary or tertiary structure. IDRs are prevalent in the human proteome,^32^ and growing evidence indicates that they have key roles in processes such as epigenetic and transcriptional regulation, signal transduction, cell cycle, and molecular condensate formation.^32–36^ IDRs often mediate protein-protein interactions and can function across a spectrum of binding modes.^32,37^ In coupled binding and folding, an IDR adopts a static conformation when bound to a structured partner. In contrast, in “fuzzy” binding, an IDR engages its partner through a dynamic, heterogeneous ensemble of conformations. Disordered complex formation can occur when both the IDR and partner remain disordered in the bound state. The primary sequence of an IDR biases its potential binding modes. The amino acid sequence determines the possibility of secondary structure formation and may encode SLIMs. In addition, the amino acid content and patterning may confer physicochemical specificity to the IDR.

Here, we investigate how the intrinsically-disordered RD3 of human PUM1 and PUM2 interacts with the CCR4–NOT complex to mediate mRNA repression. Using functional and biochemical approaches, we dissect RD3 and identify minimal peptides that retain the ability to repress mRNAs. Contrary to the hypothesis that these peptides contain SLIMs, we find that their linear peptide sequence is not critical. Instead, aliphatic and aromatic residues are the key determinants of RD3 repressive activity and its ability to recruit CCR4–NOT. Crosslinking mass spectrometry of a reconstituted PUM1 RD3:CCR4–NOT complex to identify proximal residues reveals multiple contacts between RD3 and CNOT1, CNOT2, and CNOT3. Mapping these contact regions onto a three-dimensional structure of the NOT module indicates that the crosslinks must represent an ensemble of RD3 interactions. Together, these results support a model in which the physiochemical properties of human PUM RD3 enable multivalent, fuzzy interactions with the NOT module, contributing to recruitment of CCR4-NOT and resulting in robust repression of PUM target mRNAs.

## Results

### RD3 is necessary and sufficient for full PUM repressive activity

We previously demonstrated that the repressive activity of PUM1 and PUM2 is conferred by their N-terminal regions.^28,29^ These regions, spanning amino acid residues 1-827 for PUM1 and 1-704 for PUM2, were subdivided into three repression domains (RD1-3) and two Pumilio conserved motifs (PCMa, PCMb) based on sequence conservation and function (**Figure 1A**).^6,28,29^ Prior work also showed that RD3 (**Figure 1A**. PUM1 589-827 and PUM2 471-704) exhibited robust repressive activity when targeted to a reporter mRNA in a tethered function assay.^28^ To further study the activity of RD3, we used an established dual luciferase reporter assay to measure and compare the repressive activity of full length PUM1 or PUM2 to versions lacking RD3.^7,28^ This system uses PUMs with altered RNA-binding specificity, wherein the RNA-recognition amino acid residues of the 6th repeat of the RBD are programmed to bind to a modified PRE with a 5′-UGGACAUA sequence (PRE UGG). This approach enabled us to measure the repressive activity of PUM effectors with altered specificity in wild-type cells, because the endogenous PUM1 and PUM2 are unable to bind to the modified PRE UGG.^7,28^

**Figure 1.**
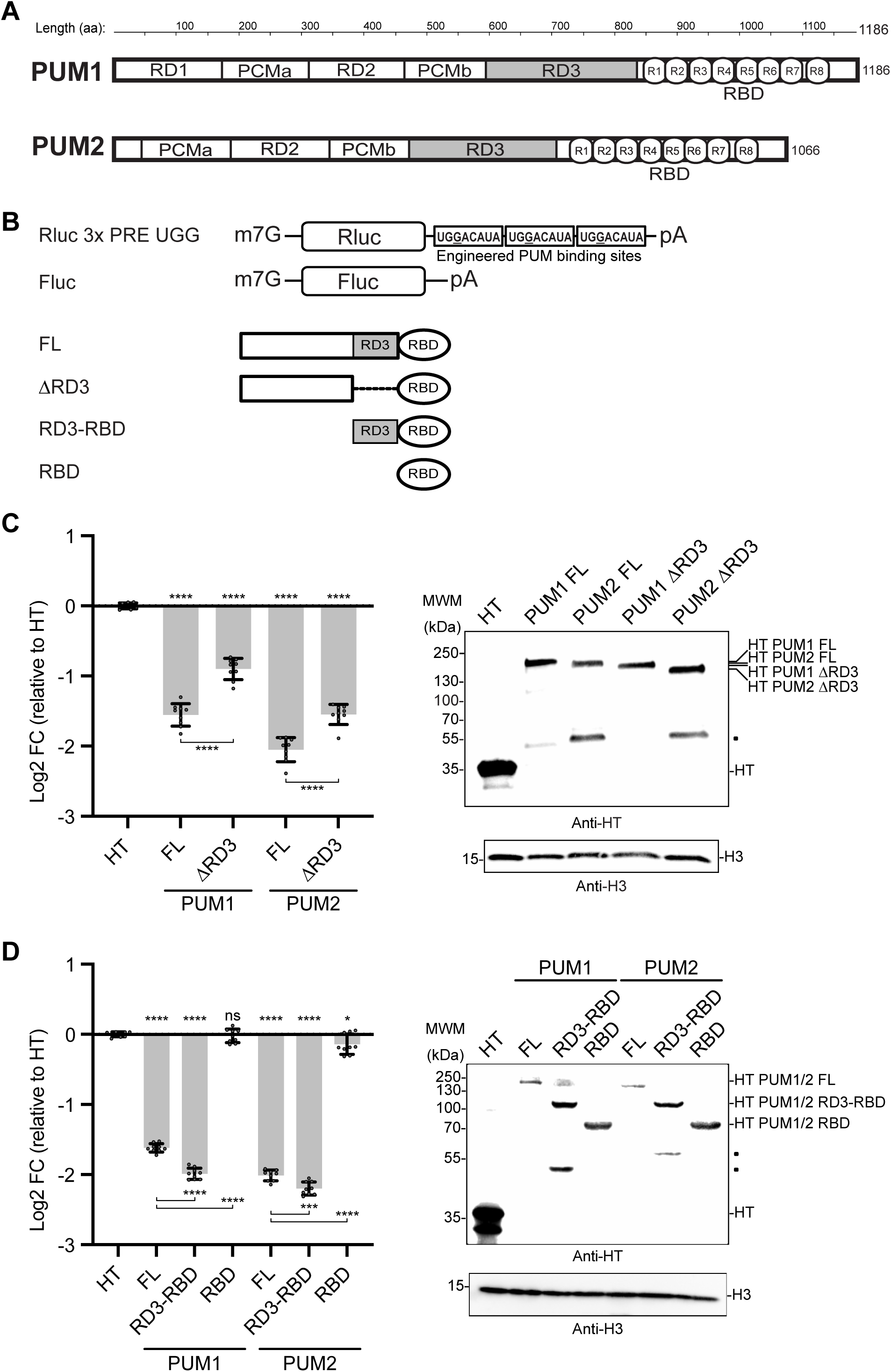
RD3 is necessary and sufficient for full PUM repressive activity. (A) Diagrams of PUM1 and PUM2 protein domain structure. PUM1 and PUM2 are drawn to scale, with repression domains (RD1-3), Pumilio conserved motifs (PCMa and PCMb), and the RNA-binding domain (RBD) with eight PUF repeats (R1-8) indicated.^6,29^ RD3, which confers the majority of repressive activity of human PUMs,^28^ is shaded in grey. Amino acid (aa) residue scale is shown across the top. (B) Diagrams of *Renilla* luciferase reporter (Rluc 3x PRE UGG) and firefly luciferase internal control (Fluc) used in assays with fusion of effector regions to PUM1 RBD with altered specificity.^7^ Diagrams of effector PUM proteins are shown below; each contains an altered specificity RBD engineered to bind to a mutated Pumilio response element (PRE) containing a UGG nucleotide sequence. Three sequential altered specificity binding sites are located in the 3′ UTR of the Rluc reporter mRNA. (C) Altered specificity reporter assay comparing the repressive activity of full-length (FL) PUM1 and PUM2 to their respective RD3 deletion (ΔRD3) effectors. N=9 (three experimental repeats, each with three biological replicates). All data are shown as mean and individual points +/- standard deviation of log2 Fold Change (log2 FC) values relative to negative control HaloTag (HT). Negative values indicate repression relative to HT. Significance indicated above the x-axis denotes comparisons to HT, while significance indicated below the x-axis denotes comparisons between specific effectors. For significance calling, ns = not significant where p ≥ 0.05, p < 0.05 = *, p < 0.01 = **, p < 0.001 = ***, p < 0.0001 = **** based on ordinary one-way ANOVA and Tukey’s test for multiple comparisons. Western blot confirming expression of HT-tagged effector proteins is shown at right. Molecular weight markers (MWM) and effector protein sizes are indicated. Dot symbols indicate cross-reactive degradation products. H3 served as a loading control. (D) Altered specificity reporter assay comparing the repressive activity of FL PUMs to their respective RD3-RBD truncation and RBD alone. To achieve similar protein expression levels across effectors, 60 ng of RD3-RBD and 30 ng of RBD plasmids were transfected. N=9 (three experimental repeats, each with three biological replicates). All data are shown as mean and individual points +/- standard deviation of log2 FC values relative to HT. Western blot confirming expression of HT-tagged effector proteins is shown at right. H3 served as a loading control.

The *Renilla* luciferase reporter mRNA with three PRE UGG sites in a minimal 3′ UTR (**Figure 1B**. Rluc 3xPRE UGG) was transfected into the human HCT116 colorectal carcinoma cell line, along with an internal control firefly luciferase (**Figure 1B**. Fluc) plasmid. The repressive activity of each PUM effector, expressed as fusions to HaloTag (HT), was measured relative to an HT negative control. While full-length (FL) PUM1 and PUM2 strongly repressed reporter protein expression, repressive activity was significantly attenuated when RD3 was deleted (ΔRD3) from either PUM (**Figure 1C**). This analysis demonstrates that RD3 is necessary for full repressive activity of PUM1 and PUM2. Since the ΔRD3 proteins are still repressive, this analysis also indicates that N-terminal regions outside RD3 contribute to PUM-mediated repression.

To determine if RD3 is sufficient for repressive activity, we again used the PRE UGG reporter to measure the repressive activity of RD3 fused to the RBD from either PUM1 or PUM2, in comparison to their respective RBDs alone (**Figure 1B**). For both PUM1 and PUM2, the RBD had no or very little repressive activity, whereas the RD3-RBD effectors exerted repressive activity slightly stronger than their corresponding FL proteins (**Figure 1D**). Thus, when directed to an mRNA via a PUM RBD, RD3 is sufficient to drive robust repression. We therefore focused on this region to identify repressive mechanisms.

### RD3 is intrinsically disordered with heterogeneous sequence conservation

The N-terminal regions of PUM1 and PUM2 are predicted to be intrinsically disordered (**Figure 2A**).^6^ Sequence analysis of PUM1 RD3 (aa 589-827) revealed IDR characteristics including above-average content of small amino acid residues (25% serine, 13% glycine, and 11% alanine) and disorder-promoting amino acid residues (8% proline and 5% glutamine), along with below-average content of hydrophobic aromatic residues (4% of phenylalanine and 3% of tyrosine) and interspersed low-complexity sequences (**Figure 2B**).^38^ RD3 is also depleted of basic and acidic residues relative to the average for the human proteome. RD3 of PUM1 and PUM2 share 73% identical residues but only 15% identical residues compared with *Drosophila* Pumilio RD3, indicating the rapidly evolving nature of these IDRs. In contrast, their RBDs are highly conserved, with human PUMs sharing 80% identical residues with *Drosophila* Pumilio.^6^ Previous work on *Drosophila* Pumilio RD3 defined four conserved regions (CR1-4) based on alignment of 82 metazoan PUM homologs, and showed that CR1 and CR4 are necessary and sufficient for *Drosophila* Pumilio repressive activity.^31^ Mapping equivalent regions onto human PUMs revealed that these regions exhibit variable amino acid conservation when aligned amongst more closely-related vertebrate homologs (**Figure 2B**). In *Drosophila* Pumilio, two conserved phenylalanine residues within CR4, corresponding to F805 and F812 in PUM1 (**Figure 2B**. red stars), are necessary for RD3 activity and binding to the Not1 subunit of the CCR4–NOT complex.^31^ Recent analysis of RNA regulatory proteins in yeast indicates that hydrophobic residues are associated with repressive activity.^39^ We noted that RD3 of human PUMs have multiple interspersed phenylalanine (F), tyrosine (Y), and leucine (L) residues, some of which are conserved (**Figure 2B**. orange diamonds). This analysis of sequence characteristics directed our investigation of the function of RD3 features in mRNA repression.

**Figure 2.**
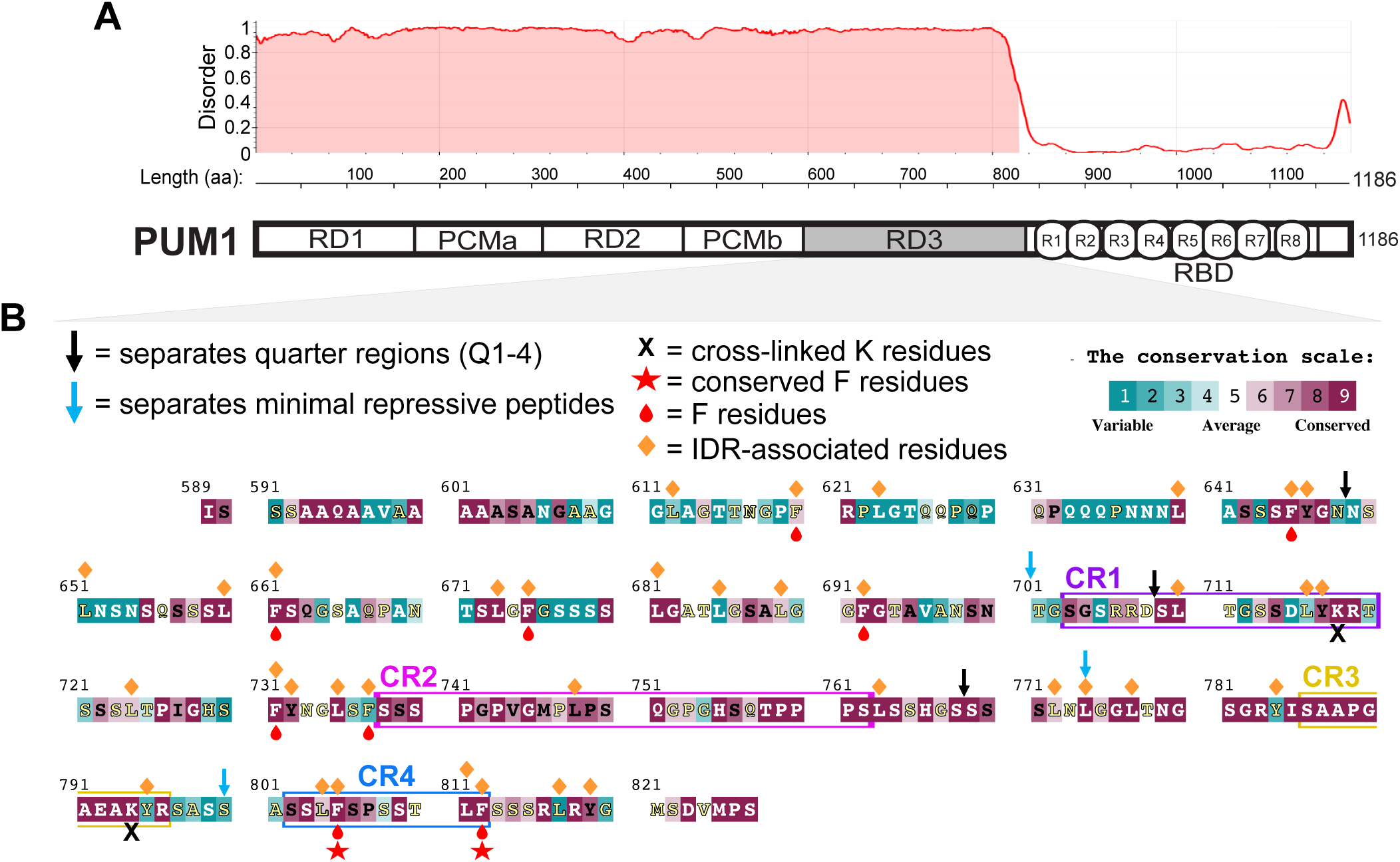
RD3 is intrinsically disordered with heterogeneous sequence conservation. (A) Diagram of PUM1 protein domain structure, RD3 is shaded in grey. Predicted protein disorder scores per amino acid residue (metapredict V3^67^) are plotted across the length of PUM1. Predicted disordered regions are indicated with red shading under the plotted line. (B) The amino acid sequence of PUM1 RD3 (aa 589-827). Sequence alignment across 74 vertebrate Pumilio homologs was performed using Clustal Omega^68^ and residues are colored according to their conservation using ConSurf^69^ scores and color scale. Conserved regions (CR1-4), defined based on sequences that are highly conserved in *Drosophila* Pumilio,^31^ are boxed and labeled in different colors. Boundaries of quarter regions of RD3 (Q1-4) are indicated with black arrows. Boundaries of regions corresponding to RD3 minimal repressive peptides are indicated with cyan arrows. Lysine (K) residues used for crosslinking experiments are indicated with black “X” symbols. Conserved phenylalanine (F) residues in CR4, necessary for RD3 function in *Drosophila* Pumilio,^31^ are indicated with red stars. All other F residues are labeled with red teardrops. Bulky/hydrophobic residues leucine (L), phenylalanine (F), and tyrosine (Y) associated with intrinsically disordered regions (IDRs) that have repressive functions^39^ are indicated with orange diamonds.

### Conserved regions of PUM1 RD3 are not necessary for repressive activity

To dissect RD3 of PUM1 and PUM2, we used an established tethered function reporter assay^28^ in which RD3 is expressed as a fusion to the RBD of the MS2 phage coat protein and targeted to a nano-luciferase reporter mRNA bearing four MS2-binding sites in the 3′ UTR (**Figure 3A**. Nluc 4x MS2 BS pA). In this context, repression depends on the tethered RD3 fragment’s ability to recruit repressive complexes to the mRNA. Expression of each RD3 effector protein was verified by detecting the V5 epitope tag included in each construct.

**Figure 3.**
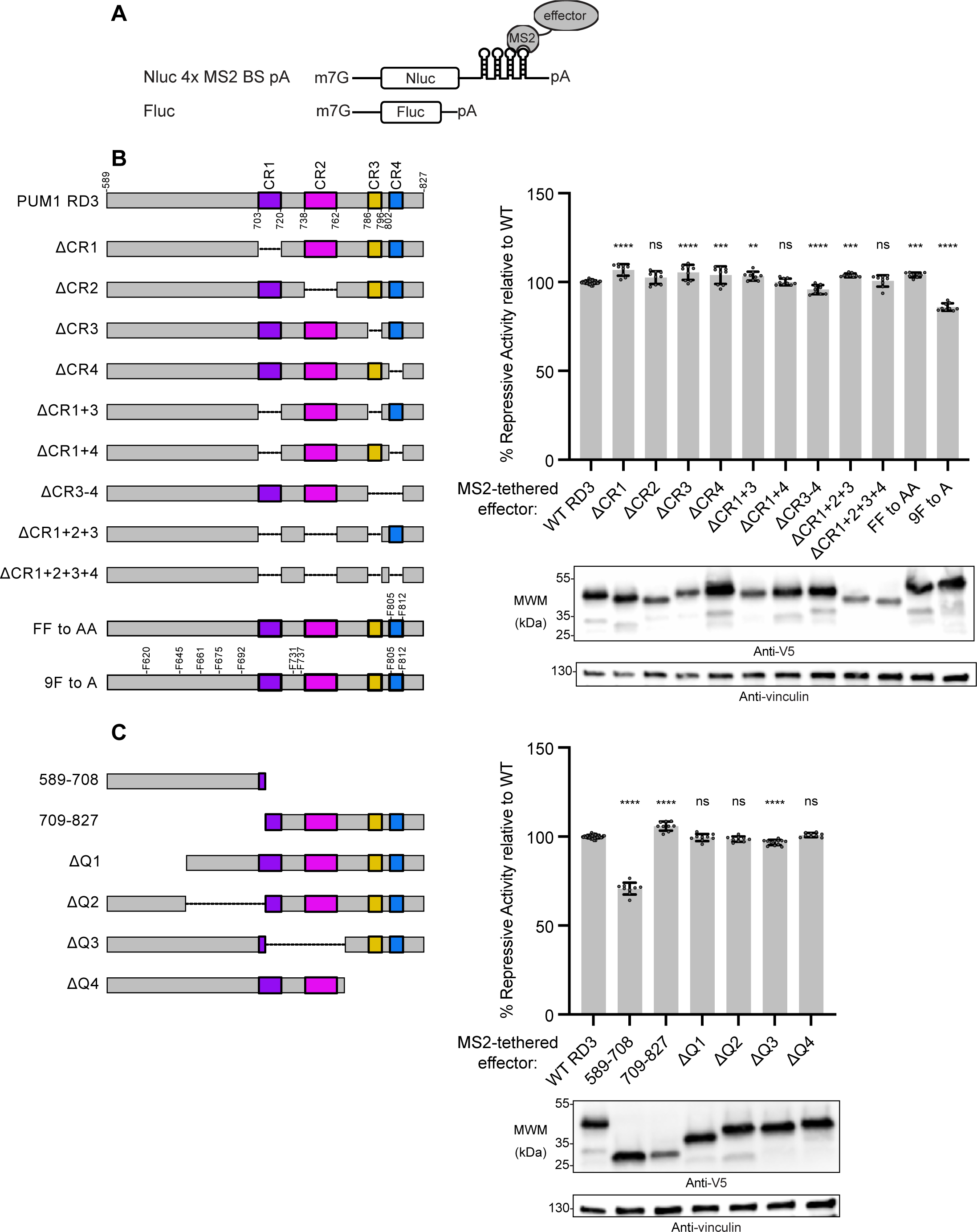
Conserved regions of PUM1 RD3 are not necessary for repressive activity; Multiple regions of RD3 are functionally redundant. (A) Diagrams of Nanoluciferase reporter (Nluc 4x MS2 BS pA) and firefly luciferase internal control (Fluc) used in tethered function assays.^28^ Four sequential MS2 binding sites located in the 3′ UTR of the Nluc reporter mRNA allow for tethering of effector proteins to the mRNA. Effector PUM proteins were expressed as fusions to MS2 coat protein and V5 epitope tag. (B) Diagram of PUM1 RD3 effector protein with conserved regions (CR1-4) and amino acid numbers denoting borders of CR1-4 indicated. Diagrams for PUM1 RD3 effectors with various CR deletions (ΔCR) are shown below. Diagrams for RD3 effectors with phenylalanine (F) residue substitutions are also shown, with relevant sites labeled. Tethered function reporter assay is shown at right comparing repressive activity of WT RD3 to RD3 effectors with individual and combined CR deletions. RD3 effectors with F805A and F812A substitutions (FF to AA) and with all F residues mutated to alanine (9F to A) were also tested. N≥9 (At least three experimental repeats, each with three biological replicates). All data are shown as mean and individual points +/- standard deviation of % Repressive Activity values relative to WT PUM1 RD3 (WT RD3). Significance indicated denotes comparisons of each effector to WT RD3. For significance calling, ns = not significant where p ≥ 0.05, p < 0.05 = *, p < 0.01 = **, p < 0.001 = ***, p < 0.0001 = **** based on ordinary one-way ANOVA and Dunnet’s test for multiple comparisons. Western blot confirming expression of V5-tagged effector proteins is shown below reporter assay. Vinculin served as a loading control. (C) Diagrams of PUM1 RD3 truncation and quarter region deletion (ΔQ1-4) effector proteins. Tethered function reporter assay comparing the repressive activity of WT RD3 to truncated RD3 effectors and quarter deletions. N≥9 (At least three experimental repeats, each with three biological replicates). All data are shown as mean and individual points +/- standard deviation of % Repressive Activity values relative to WT RD3. Western blot confirming expression of V5-tagged effector proteins is shown below. Vinculin served as a loading control.

We first examined the roles of the conserved regions by measuring repressive activity of MS2-tethered PUM1 RD3 with each CR deleted individually or in combination with others (**Figure 3B**). Neither deletion of individual nor multiple CRs impaired repressive activity relative to wild-type (WT) RD3. In fact, PUM1 RD3 with all four CRs deleted (ΔCR1+2+3+4) displayed repressive activity indistinguishable from WT (**Figure 3B**). Therefore, human PUM CRs are not required for repressive activity. This contrasts with *Drosophila* Pumilio, where deletion of CR1, CR3, or CR4 reduced RD3 activity.^31^

We also tested the importance of the two conserved phenylalanine residues that are important for *Drosophila* Pumilio RD3 activity.^31^ Substitution of F805 and F812 with alanine had no effect on RD3 repressive activity (**Figure 3B**. FF to AA). We speculated that other phenylalanine residues could functionally substitute. Therefore, we mutated all nine phenylalanine residues distributed throughout PUM1 RD3, which resulted in ∼20% reduction in repressive activity, a small, but statistically significant effect (**Figure 3B**. 9F to A). These observations were recapitulated in assays testing corresponding deletions or substitutions in PUM2 RD3 (**Figure S2A**). Taken together, the data indicate that CRs and residues important for *Drosophila* Pumilio are dispensable for human RD3 repressive activity. Although the function of RD3 and its interaction with the CCR4–NOT complex is conserved across species, the exact amino acid sequences responsible for repressive function appear to have been rewired during evolution.

### Multiple regions of RD3 are functionally redundant

We next took a systematic approach to identify functionally important sequences in RD3. We divided PUM1 RD3 into two halves. The first half of RD3 (aa 589-708) displayed repressive activity but it was diminished ∼25% relative to WT. In contrast, the second half of RD3 (aa 709-827) retained the same level of repressive activity as WT (**Figure 3C**). Since the four CRs are present in the C-terminal portion of RD3, we adjusted the N-terminal boundary of the second half to ensure CR1 was fully encompassed (**Figure S1A,** aa 701-827) and again observed repressive activity comparable to WT RD3 (**Figure S1B**). We further divided RD3 into quarters and examined the effect of deletions of individual quarters (ΔQ1-4). No deletion had a substantial effect on repressive activity (**Figure 3C**). These observations were recapitulated in our analysis of PUM2 RD3 (**Figure S2B**). Together, these results demonstrate that RD3 contains multiple repressive regions that can act in a partially redundant manner.

### Identification of minimal repressive peptides in RD3

We sought to identify minimal peptides of RD3 that retain repressive activity. We focused on PUM1 because PUM1 and PUM2 RD3 displayed similar activity patterns in our assays. As PUM1 701-827 exhibited 100% of WT RD3 repressive activity, we dissected this region into three peptides (**Figure 4A**, aa 701-774, 774-800, and 800-827). Each individual peptide possessed repressive activity, but the level was reduced relative to WT RD3 (**Figure 4B**). We also measured the contribution of each peptide by deleting the region in the context of PUM1 701-RBD, using the Rluc 3xPRE UGG reporter (**Figure 4A**). If the repressive activity is additive, we expected that all deletions would reduce repression. Deletion of 701-774 (774-RBD) reduced repressive activity, but deletion of 774-800 (701-RBD Δ774-800) or 800-827 (701-RBD Δ800-827) had no effect, relative to the parental 701-RBD effector protein (**Figure 4C**). These data indicate that the repressive function of the minimal RD3 peptides is predominantly redundant, not additive.

**Figure 4.**
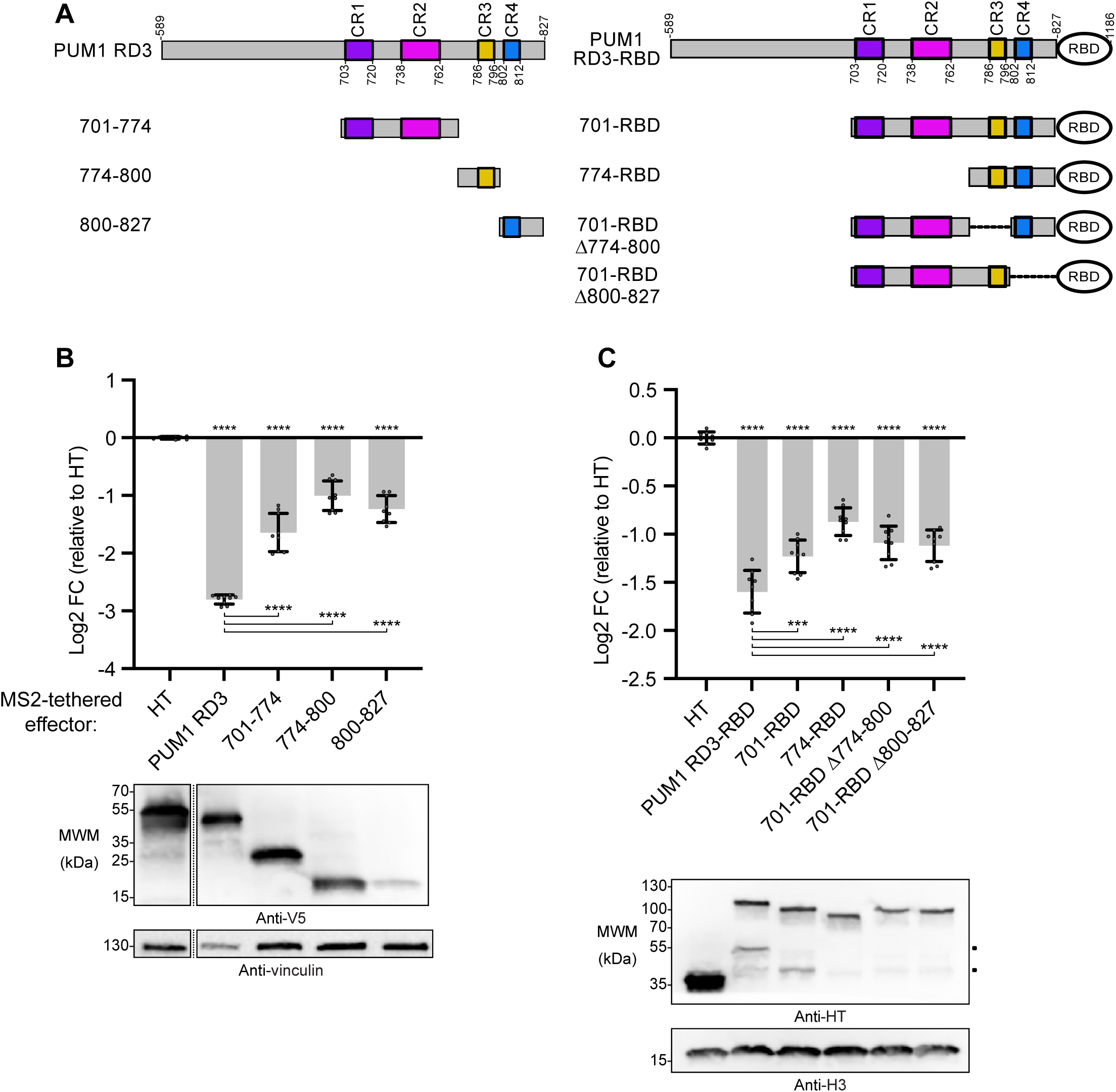
Identification of minimal repressive peptides in RD3. (A) Diagram of minimal PUM1 RD3 and RD3-RBD effectors with amino acid numbers indicated. Effectors fused to MS2 coat protein are shown at left, while effectors fused to PUM1 RBD with altered specificity are shown to the right. (B) Tethered function reporter assay comparing repressive activity of WT RD3 to minimal RD3 region effectors. N=9 (three experimental repeats, each with three biological replicates). All data are shown as mean and individual points +/- standard deviation of log2 Fold Change (log2 FC) values relative to negative control HaloTag (HT). Significance indicated above the x-axis denotes comparisons to HT, while significance indicated below the x-axis denotes comparisons between specific effectors. For significance calling, ns = not significant where p ≥ 0.05, p < 0.05 = *, p < 0.01 = **, p < 0.001 = ***, p < 0.0001 = **** based on ordinary one-way ANOVA and Tukey’s test for multiple comparisons. Western blot confirming expression of V5-tagged effector proteins is shown below. Vinculin served as a loading control. Dashed vertical lines indicate that blot images were cropped to show relevant lanes. (C) Altered specificity reporter assay comparing the repressive activity of PUM1 RD3-RBD to effectors with combinations of RD3 minimal regions fused to PUM1 RBD with altered specificity. For each PUM effector, 60 ng of effector plasmid was transfected. N=9 (three experimental repeats, each with three biological replicates). All data are shown as mean and individual points +/- standard deviation of log2 FC values relative to HT. Western blot confirming expression of HT-tagged effector proteins is shown below. H3 served as a loading control.

### RD3 contains multiple repressive peptides that contact CCR4–NOT

The PUM N-terminus interacts with the CCR4–NOT deadenylase complex primarily through contacts between RD3 and the NOT module containing the CNOT1, CNOT2, and CNOT3 subunits (**Figure 5A**).^28^ Using our functional data as a guide, we sought to identify which regions of RD3 promote interaction with the NOT module. To do this, we performed *in vitro* pull-down assays using recombinant, purified proteins. In these experiments, we used various StrepII affinity-tagged maltose binding protein (MBP)-PUM1 RD3 constructs as baits (**Figure 5B-C**). We observed that the PUM1 701-RBD protein containing the minimal repressive peptides interacted with the NOT module (**Figure 5C**). Neither MBP nor the PUM1 RBD alone bound to the NOT module, whereas PUM1 RD3-RBD with the complete RD3 did bind, demonstrating specificity of the CCR4–NOT interaction with RD3 in these assays (**Figure 5D**). Binding to the NOT module was also detected for 774-RBD and RD3-RBD Δ774-827 constructs (**Figure 5D**). Additionally, we tested NOT module binding with the minimal RD3 repressive peptides. Using 589-701 (**Figure 5E**), 701-774 (**Figure 5F**), 774-800 (**Figure 5G**), and 800-RBD (**Figure 5G**) as baits in pull-down assays, we detected binding to the NOT module for each construct. These results indicate that there are at least four independent contacts linking RD3 to the NOT module (summarized in **Figure 5B**). Moreover, these contact regions correspond to those exhibiting repressive activity in functional assays.

**Figure 5.**
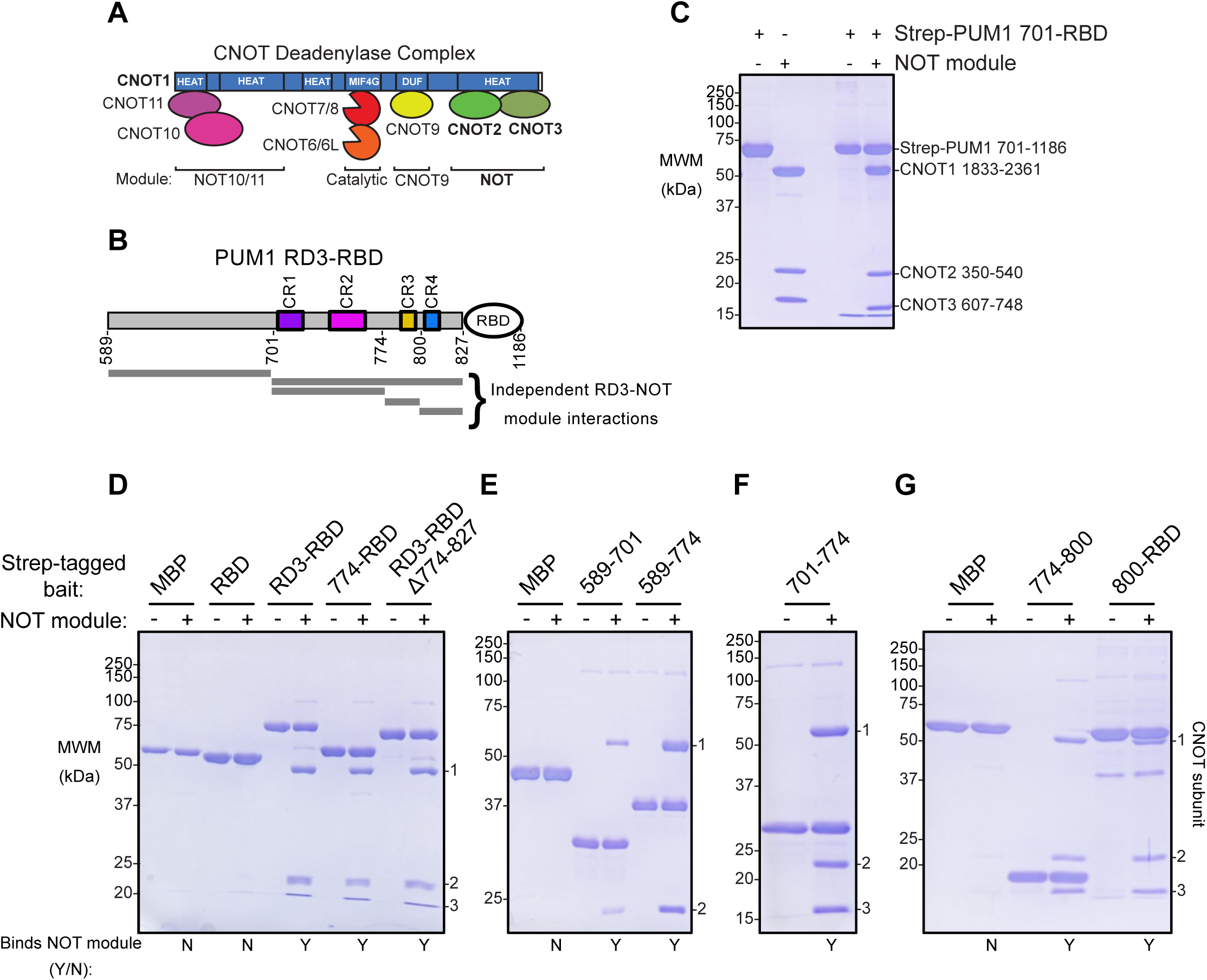
RD3 contains multiple repressive peptides that contact CCR4–NOT. (A) Diagram of the CCR4–NOT deadenylase complex architecture. Subunits are grouped by the indicated functional modules. (B) Diagram of PUM1 RD3-RBD protein with amino acid residue numbers labeled. Grey bars summarizing regions that engage in independent interactions with the NOT module are shown below. (C) SDS-PAGE and Coomassie blue staining of recombinant, purified StrepII-tagged MBP fusion PUM1 701-1186 and NOT module proteins (input, left lanes) and detected in an *in vitro* pull-down of StrepII-tagged MBP-PUM1 701-1186 (right lanes). (D-G) SDS-PAGE and Coomassie blue staining of *in vitro* pull-downs with indicated recombinant, purified StrepII-MBP-PUM1 baits. NOT module subunits (1-3) detected in pull-downs are labeled to the right of each gel image. Binary assessment of binding to the NOT module (Yes/No) is indicated below each pull-down. StrepII-MBP and StrepII-MBP-PUM1 RBD served as negative controls for NOT module interaction.

### Crosslinking mass spectrometry reveals a fuzzy binding mode of RD3:NOT module interaction

Consistent with the role of RD3 in NOT module interaction and repressive activity, we found that purified PUM1 RD3-RBD formed a stable complex with the NOT module, co-eluting by size-exclusion chromatography, while the RBD alone did not (**Figure S3A-B**). Despite purification of this stable complex, attempts to determine a high-resolution structure of RD3 peptide bound to the structured NOT module to identify specific points of contact were unsuccessful. IDRs such as RD3 present a challenge for structural determination, as X-ray crystallographic and high resolution single-particle cryo-electron microscopy approaches rely on well-ordered and homogenous complexes, whereas IDRs introduce flexibility and multiple contact points add heterogeneity.

As an alternative approach to characterize the PUM1 RD3:NOT module interaction, we performed crosslinking mass spectrometry to identify neighboring regions of RD3 and CCR4– NOT. Recombinant PUM1 RD3-RBD (aa 589-1186) was incubated with a PRE-containing RNA ligand and then recombinant NOT module was added. The resulting ribonucleoprotein complex was crosslinked using bis(sulfosuccinimidyl)suberate (BS3), an amine-to-amine crosslinker with a spacer length of 11.4 Å. We observed a band of ∼170 kDa on an SDS-PAGE gel, corresponding to the crosslinked PUM1:NOT complex (**Figure S3C**). Two technical replicate samples were digested with a trypsin/LysC mix, and fragments were analyzed by mass spectrometry to identify crosslinked residues. By plotting the inter-protein crosslinks on the protein sequences, we observed that the two lysine residues in PUM1 RD3 (K718 and K794) crosslinked to multiple lysine residues in different CCR4–NOT subunits (**Figure 6A**). RD3 K718 crosslinked to multiple sites on the CNOT1, CNOT2, and CNOT3 subunits, and RD3 K794 crosslinked to CNOT1 and CNOT2 subunits (**Figure 6A-C**).

**Figure 6.**
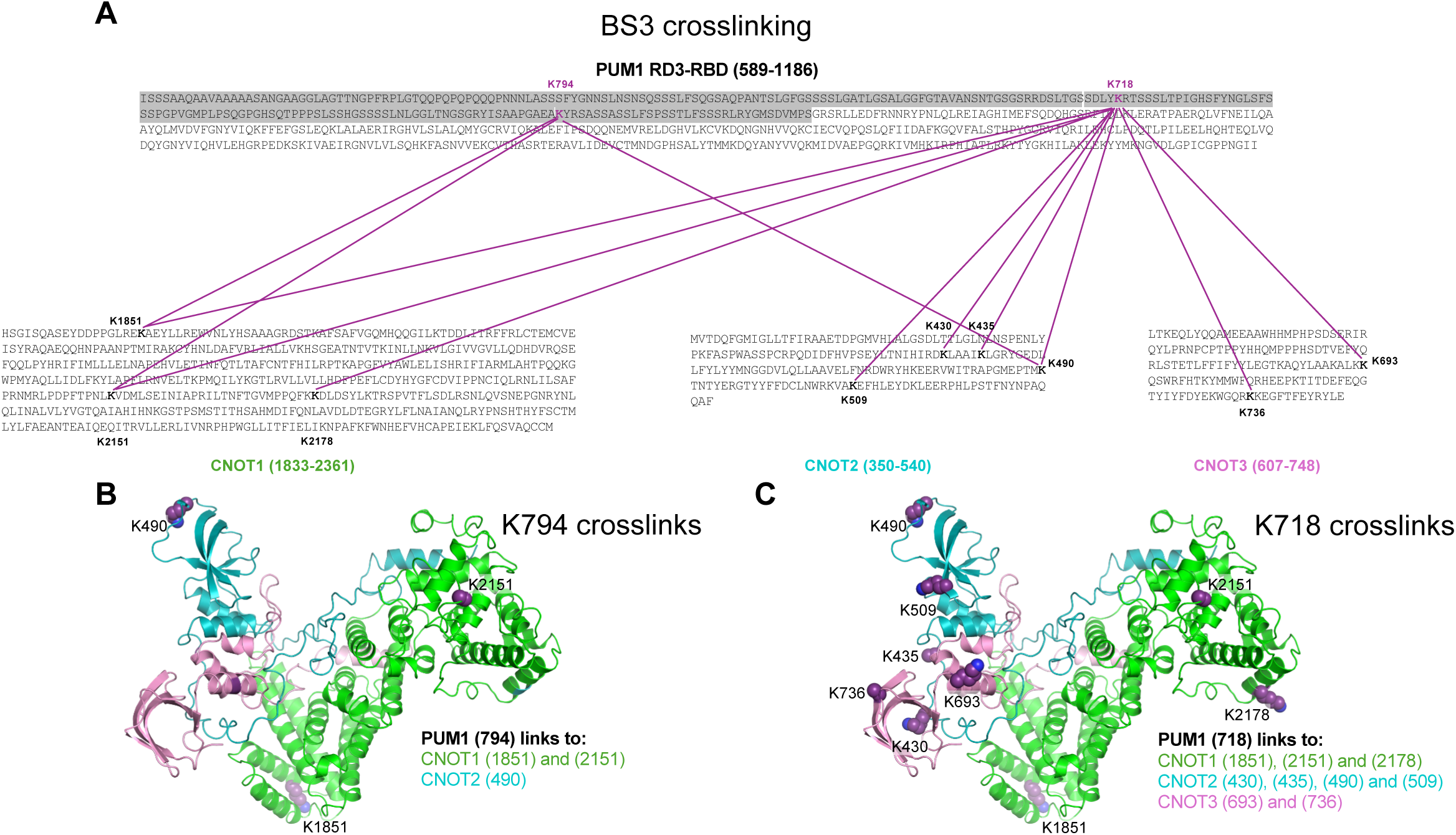
Crosslinking mass spectrometry reveals a fuzzy binding mode of RD3:NOT module interaction. (A) Amino acid sequence of PUM1 RD3-RBD. RD3 is highlighted in grey and lysine (K) residues available for BS3 crosslinking are magenta. Amino acid sequences for the regions of CNOT1, CNOT2, and CNOT3 used to form the NOT module are shown below. Crosslinked residues in each subunit are bolded, with magenta lines indicating the locations of inter-protein crosslinks found in the RD3-RBD:NOT module complex. (B) Crystal structure of the NOT module with residues that crosslink to PUM1 K794 shown with purple spheres and labeled. (C) Same as in (B) with residues that crosslink to PUM1 K718 shown with purple spheres and labeled.

We visualized the distribution of crosslink sites on the crystal structure of the NOT module (PDB 4C0D)^40^ (**Figure 6B-C**). Rather than observing a distinct region of crosslinking for each PUM1 RD3 lysine residue, we found that each residue crosslinked to multiple “hotspot” regions (**Figure 6B-C**). The crosslink hotspots for each PUM1 RD3 lysine were spatially distant and must represent independent, mutually exclusive binding events. This phenomenon indicates that crosslinks did not form from a single distinct conformation. Instead, RD3 is flexible and dynamically interacts with the NOT module subunits through an ensemble of conformations.

### The activity of RD3 repressive peptides withstands targeted deletions or substitutions

Both the RD3 structure-function analysis and crosslinking experiments presented above support the model that RD3 makes redundant multivalent contacts with CCR4–NOT to repress mRNAs. We hypothesized that the minimal RD3 repressive peptides may each contain conserved SLIMs that mediate CCR4–NOT recruitment and repressive activity. We therefore attempted to identify functional motifs by making further deletions/substitutions within the minimal peptides (**Figure S4**). For the 701-774 repressive peptide, deletion of CR1, CR2, or CR1+2 together caused minor reductions in repressive activity in the tethered function assay. For the 774-800 repressive peptide, we mutated a conserved proline (P789). The P789A substitution reduced, but did not eliminate, repressive activity of 774-800 (**Figure S4**). Mutation of the two conserved phenylalanine residues (F805 and F812) in CR4^31^ also decreased, but did not eliminate, the repressive activity of the 800-827 effector (**Figure S4**). The residual repressive activity despite targeted deletions/substitutions demonstrated that evolutionary conservation was insufficient to elucidate the sequences or properties necessary for RD3 activity. We therefore pursued alternative means to assess whether the linear polypeptide sequence of RD3 was essential.

### Shuffling amino acid sequences does not perturb the activity of RD3 repressive peptides

Recent insights into the properties of IDRs provided us with a framework for examining how their sequences may contribute to their molecular functions.^32,33,41^ IDRs may rely on specific linear sequence motifs (e.g. SLIMs) but they may also rely on the physicochemical properties conferred by the amino acid composition.^42^ To distinguish between these mechanisms for RD3, we randomly shuffled the positions of amino acid residues within each peptide while maintaining the same composition (**Figure 7A**). We anticipated that shuffling would inactivate SLIM-dependent repressive activity, but would not affect repression if it was reliant on the unaltered physicochemical properties. Each shuffled peptide revealed repressive activity that was indistinguishable (701-774 and 774-800) or increased (800-827) relative to its non-shuffled counterpart (**Figure 7B**). These results indicate that the amino acid residue content, and thus physicochemical composition, confers repressive activity of RD3 peptides, rather than specific linear sequence.

**Figure 7.**
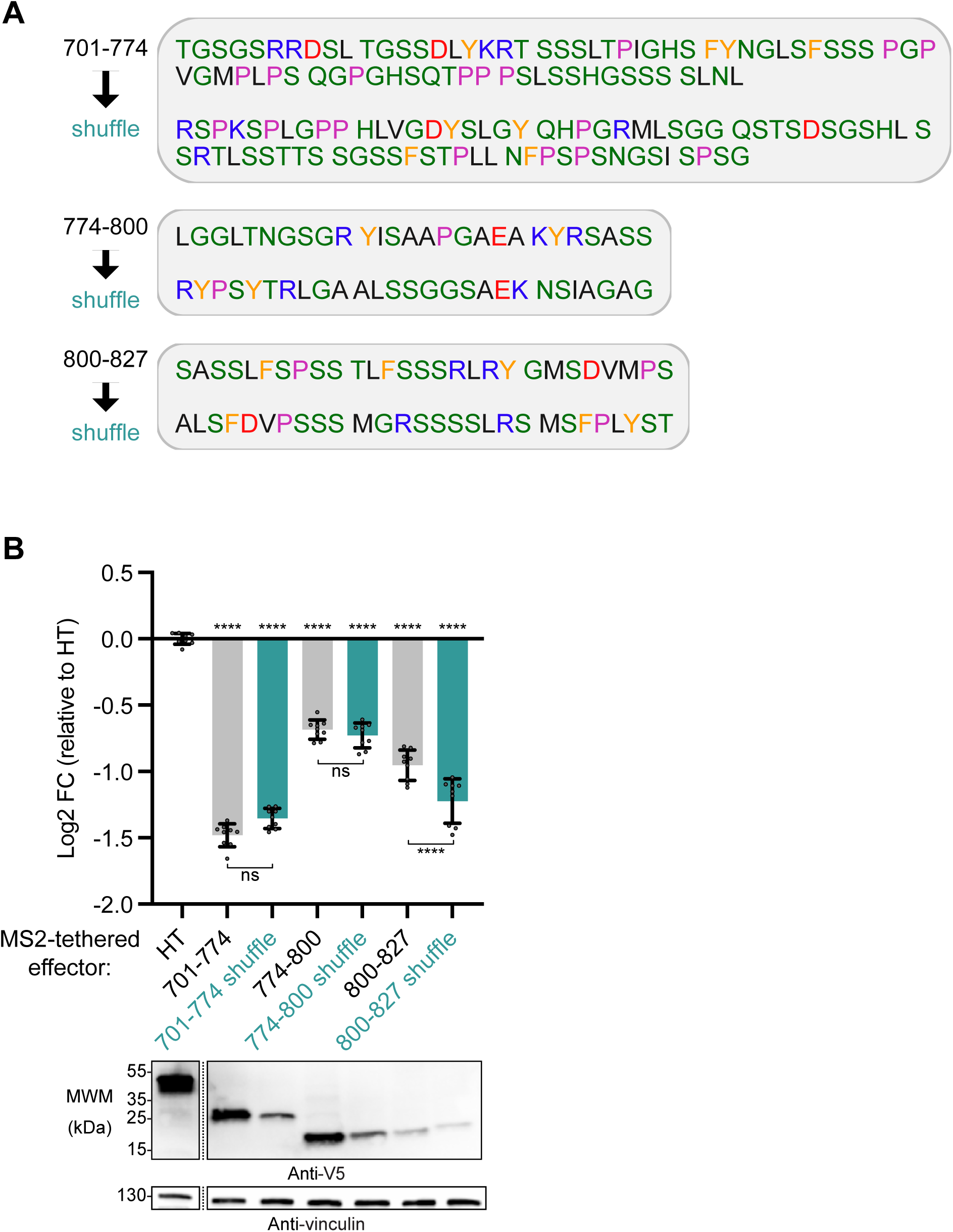
Shuffling amino acid sequences does not perturb the activity of RD3 repressive peptides. (A) Amino acid sequences of PUM1 RD3 minimal repressive peptides and their corresponding effectors with randomly-shuffled amino acid residues. Residues are colored based on their biochemical properties using the CIDER color scheme (black=aliphatic, green=polar, blue=basic/positive, red=acidic/negative, pink=proline, yellow=aromatic). (B) Tethered function reporter assay comparing repressive activity of each minimal RD3 peptide to its shuffled counterpart. N=9 (three experimental repeats, each with three biological replicates). All data are shown as mean and individual points +/- standard deviation of log_2_ Fold Change (log2 FC) values relative to negative control HaloTag (HT). Significance indicated above the x-axis denotes comparisons to HT, while significance indicated below the x-axis denotes comparisons between specific effectors. For significance calling, ns = not significant where p ≥ 0.05, p < 0.05 = *, p < 0.01 = **, p < 0.001 = ***, p < 0.0001 = **** based on ordinary one-way ANOVA and Tukey’s test for multiple comparisons. Western blot confirming expression of V5-tagged effector proteins is shown below. Vinculin served as a loading control.

### Aliphatic and aromatic amino acid residues are essential for RD3 activity

A recent analysis of IDRs in regulatory proteins of *Saccharomyces cerevisiae* identified hydrophobic and aromatic amino acid residues as important determinants of function.^39^ That observation inspired us to test the necessity of the 36 lysine (L), phenylalanine (F), and tyrosine (Y) residues that are interspersed throughout RD3 for repressive function (**Figure 8A**).

**Figure 8.**
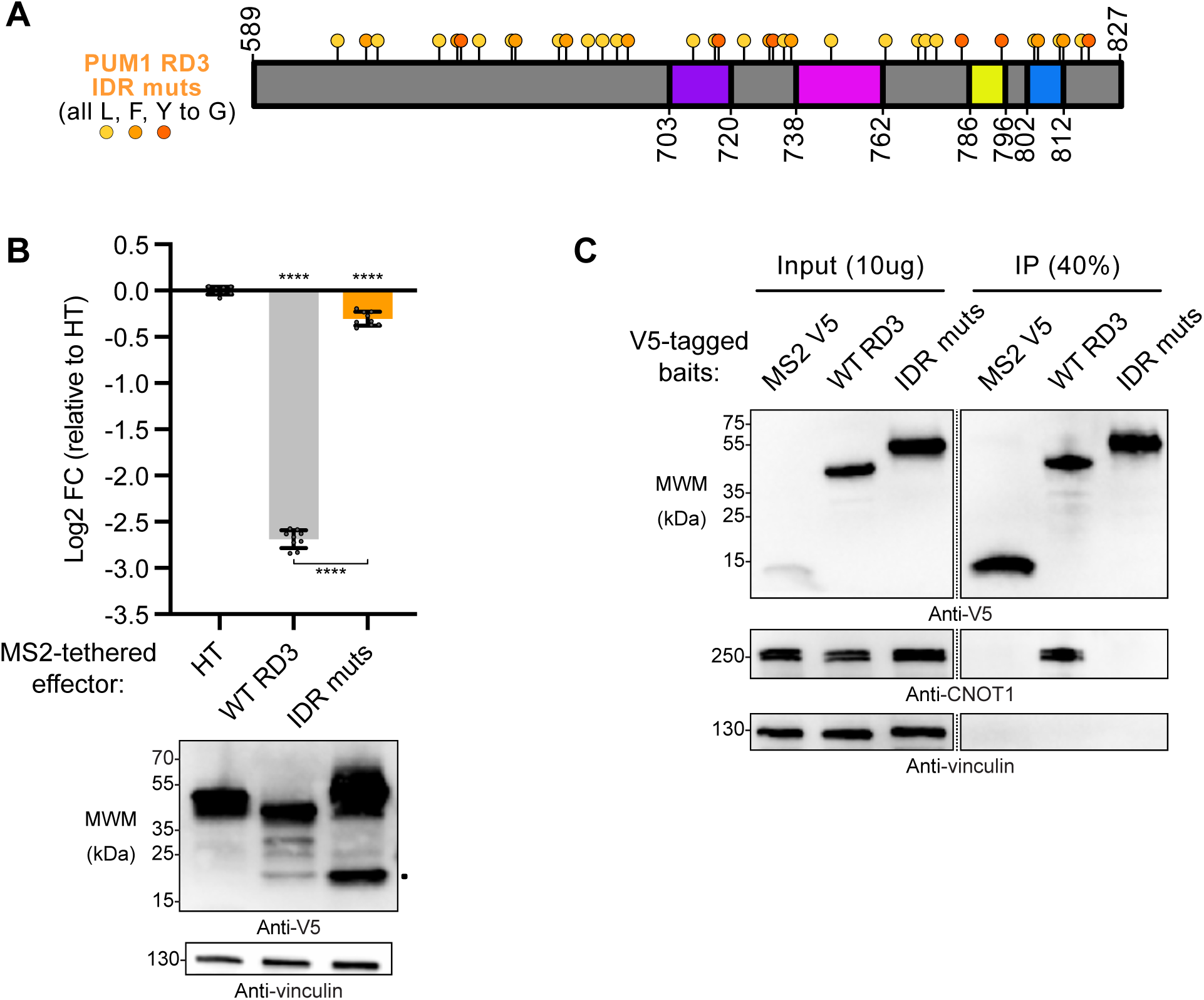
Aliphatic and aromatic amino acid residues are essential for RD3 activity. (A) Diagram of PUM1 RD3 effector protein with all IDR-associated residues^39^ L, F, and Y mutated to G (IDR muts). Position and identity of amino acid residue in the WT sequence are indicated. Tethered function reporter assay comparing repressive activity of WT RD3 to the IDR muts effector. N=9 (three experimental repeats, each with three biological replicates). All data are shown as mean and individual points +/- standard deviation of log2 Fold Change (log2 FC) values relative to negative control HaloTag (HT). Significance indicated above the x-axis denotes comparisons to HT, while significance indicated below the x-axis denotes comparisons between specific effectors. For significance calling, ns = not significant where p ≥ 0.05, p < 0.05 = *, p < 0.01 = **, p < 0.001 = ***, p < 0.0001 = **** based on ordinary one-way ANOVA and Tukey’s test for multiple comparisons. Western blot confirming expression of V5-tagged effector proteins is shown below. Vinculin served as a loading control. (C) Co-immunoprecipitation of endogenous CNOT1 from HCT116 cell lysate by V5-tagged WT and IDR muts RD3. MS2-V5 served as a negative control bait protein. Vinculin served as a negative control for non-specific binding.

Compared to WT PUM1 RD3, substitution of all 36 L, F, and Y residues with glycine (IDR muts) nearly eliminated repression in the tethered function assay (**Figure 8B**). We then tested the effect of these substitutions on interaction with CCR4–NOT by performing co-immunoprecipitation analysis using transfected WT or IDR mut versions of PUM1 RD3 with a V5 epitope tag. We immunoprecipitated tagged proteins from cell extract with V5 antibody-conjugated beads and then samples probed by western blot for the presence of endogenous CNOT1, the central scaffold protein of the CCR4–NOT complex. CNOT1 co-immunoprecipitated with WT RD3 but not the IDR muts effector or the V5-tagged MS2 coat protein negative control (**Figure 8C**). We concluded that the L, F, and Y residues throughout RD3 are essential for both repressive activity and the ability to interact with CCR4–NOT, consistent with the model that RD3 contributes to PUM repression by recruiting the deadenylase complex.

## Discussion

RD3 of human PUM1 and PUM2 illustrates how intrinsically disordered regions can recruit CCR4–NOT through composition-driven, fuzzy binding to control mRNA decay. Rather than acting through a single short linear motif, RD3 functions as a multivalent interaction platform whose repressive activity is distributed across the domain (**Figure 9**). Our data point to the amino acid composition and the distribution of aliphatic and aromatic residues, rather than the primary sequence, as central to this mechanism. In this view, CCR4–NOT recruitment by PUM RD3 reflects a multivalent and dynamically heterogeneous engagement of the NOT module. We propose that PUM-mediated repression is a model for how disordered effector domains achieve robust yet flexible regulation of mRNA turnover.

**Figure 9.**
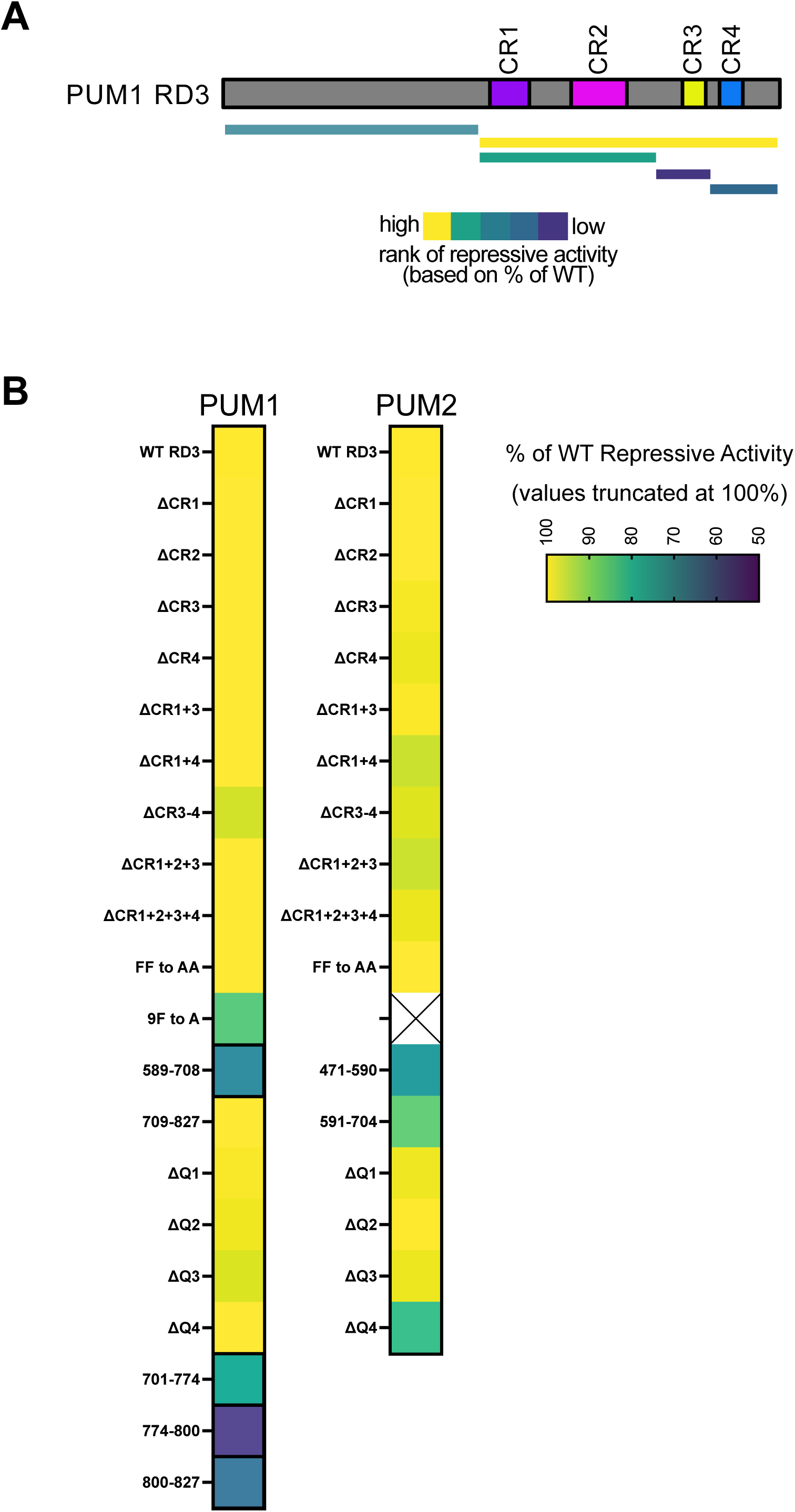
Four minimal regions of PUM1 RD3 can independently contact CCR4–NOT to drive PUM-mediated repression of mRNAs. (A) Summary of regions of PUM1 RD3 with independent repressive activity that contact the NOT module. Each region is shown as a bar below the PUM1 RD3 protein diagram and is colored according to its rank of repressive activity relative to the other interacting regions. The bar for 701-827 encompassing CR1-4, while not indicating a minimal repressive peptide, is shown as a reference for an effector capable of full WT RD3 repressive activity. (B) Heatmaps representing % of WT Repressive Activity values for each PUM1 and PUM2 construct tested in Figure 3 **and Figure S2**. Any effectors with higher activity than WT are displayed at the maximum value of 100%. Box outlines are bolded for minimal repressive peptides.

The same composition-based features that make RD3 a fuzzy interaction platform also make it modular. We identified four minimal repressive peptides in RD3 (summarized in **Figure 9**) that can independently repress and interact with CCR4–NOT. Identification of crosslinks between RD3 and the NOT module indicate proximity of specific RD3 residues to multiple regions of the three subunits: CNOT1, CNOT2, and CNOT3. We demonstrated that combinations of multiple minimal RD3 peptides also retain strong repressive activity when fused to the PUM RNA-binding domain, which suggests that these short segments can be treated as portable effector modules. Coupling RD3 peptides to programmable PUM-based^43–46^ or heterologous RNA-binding domains could, in principle, generate synthetic repressors that recruit CCR4–NOT to defined mRNA targets. Because CCR4–NOT is broadly expressed, such constructs might be applied in many cell types, provided that their potency and specificity are tuned carefully.

At the same time, RD3 is not the sole PUM effector.^28^ While RD3 provides the dominant CCR4– NOT interaction surface, other N-terminal regions contribute additional, weaker repression (**Figure 1C**). This layered architecture suggests that PUM integrates several disordered segments to fine-tune mRNA decay. Identifying these auxiliary regions, defining their binding partners, and testing whether they engage distinct CCR4–NOT interfaces or alternative decay pathways will be essential for a complete mechanistic picture of PUM-mediated regulation.

The intrinsically-disordered nature of RD3 enables it to bind the three subunits of the NOT module in a dynamic ensemble of conformations, using multiple points of contact dependent on the aliphatic and aromatic residues. IDRs are uniquely poised to participate in fuzzy-binding interactions, as they are rapidly-evolving and often use multiple contacts to build in redundancy, rendering them more resilient to mutations that might disrupt the interaction interface.^32^ Moreover, fuzzy binding allows for dynamic, reversible interactions, which may be important during responses to stimuli or developmental transitions.^34,41^ For PUMs and other CCR4–NOT interactors, multivalent contacts likely provide affinity and specificity for the interaction while allowing for dynamic, transient recruitment of CCR4–NOT to degrade target mRNAs and tightly control gene expression.

Growing evidence indicates that PUM proteins from yeast to mammals use IDRs to recruit CCR4–NOT.^7,28–31,39,47–49^ In *Drosophila* Pumilio, three N-terminal repression domains each bind CCR4–NOT and contribute to repression, and RD3 uses conserved regions to contact central CNOT1 and the NOT module.^29–31^ In fission yeast, an IDR in Puf3 forms redundant contacts with CNOT9, CNOT1, CNOT2, and CNOT3, with tryptophans in at least two SLIMs contributing to binding.^48^ Repressive IDRs have also been identified in budding yeast Mpt5/Puf5,^39^ which represses mRNAs via CCR4–NOT and decapping, though its binding determinants are less defined.^39,47,50,51^ Despite this shared mechanism, these IDRs show little primary sequence homology to metazoan Pumilio proteins, underscoring the plasticity of IDR-mediated interactions.^6,32^

Other RNA-binding repressors utilize multivalent IDRs to recruit CCR4–NOT.^52^ TTP, which binds to AU-rich elements in cytokine mRNAs, engages three CCR4–NOT interfaces through IDRs that contact CNOT9, central CNOT1, and the NOT module.^53,54^ GW182/TNRC6 proteins that are necessary for miRNA-mediated repression use multiple tryptophan residues that dock into hydrophobic pockets on CNOT9 and the NOT module.^55–57^ Vertebrate Nanos uses an aromatic SLIM that binds a hydrophobic patch on CNOT1,^58^ whereas *Drosophila* Nanos uses two helical SLIMs to interact with CNOT1 and CNOT3.^59^ These examples initially suggested that human PUM RD3 would depend on a small number of rigid SLIMs.^52^ Instead, our dissection shows that RD3 activity is mainly composition- and hydrophobicity-driven and tolerates extensive sequence shuffling.

Overall, these systems highlight a recurring role for aliphatic and aromatic residues at CCR4– NOT interfaces, where hydrophobic and π interactions engage both structured domains and disordered segments.^32,52^ For PUM RD3, leucine, phenylalanine, and tyrosine residues are required for function, but how interchangeable they are and how many, in which patterns, define a minimal module remains unclear. More broadly, IDR functions remain challenging to predict from sequence alone, because many regions, including RD3, depend on composite properties that sequence conservation captures poorly ^32,60^ Our results argue that composition, residue patterning, and protein context must be considered, and that sequence shuffling is a practical way to distinguish SLIM-driven from composition-driven interactions. Loss of function after shuffling points to critical motifs or patterns, whereas retained activity, as for RD3, indicates a dominant role for composition and physicochemical properties, with both mechanisms likely coexisting in many IDRs^32,42^

Taken together, the functional redundancy of RD3 peptides, their tolerance to sequence shuffling, their reliance on hydrophobic and aromatic residues, and crosslinking data revealing multiple mutually exclusive contacts all support a fuzzy ensemble binding mode in which multivalent, composition-driven interactions enable robust yet reversible recruitment of CCR4– NOT to PUM target mRNAs.

### Experimental procedures

#### Plasmids and cloning

All plasmids used in this study are listed in **Supplemental Table S1**. The nanoluciferase (Nluc) reporter plasmid for tethered function experiments (pNLP 4x MS2 BS pA) and *Renilla* luciferase (Rluc) reporter plasmid for altered specificity experiments (psiCheck 3x UGG) were previously described.^28^ For all experiments, the firefly luciferase (Fluc) plasmid (pGL4.13) (Promega) was co-transfected as a control for transfection efficiency. Plasmids for expression of PUM tethered function effector proteins were cloned as previously described, with effector proteins expressed from pF5K vector as fusions to an N-terminal MS2 coat protein (MS2CP) and a V5 epitope tag.^28,61^ Deletion and truncation constructs were cloned using inverse PCR, while single and double amino acid mutations were cloned using site-directed mutagenesis (Agilent QuikChange II). The tethered PUM1 RD3 9F to A and IDR muts constructs were created by restriction cloning gene fragments containing the mutations (IDT or Twist Bioscience) into the pF5K MS2CP V5 backbone. Shuffled constructs were designed by randomly shuffling the codons in each region, using atmospheric noise as a randomness generator algorithm (https://www.random.org/lists/), and then gene fragments containing the shuffled sequences (IDT) were cloned into the pF5K MS2CP V5 vector. Plasmids for expression of altered specificity PUM effector proteins were cloned as previously described, with effector proteins containing the altered specificity PUM RNA-binding domain (R6as) expressed from pFN21A vector as fusions to an N-terminal HaloTag.^7,28,62^ Deletion and truncation constructs were cloned using inverse PCR.

#### Cell culture

HCT116 cells (ATCC) were incubated at 37 ℃ with 5% CO2 and cultured in McCoy’s 5A media (Thermo Fisher Scientific) supplemented with 10% fetal bovine serum (Genesee or Sigma-Aldrich) and 1x antibiotics containing 100 U/mL penicillin and 100 µg/mL streptomycin (Thermo Fisher Scientific).

#### Luciferase reporter assays

For luciferase reporter assays, 5,000 HCT116 cells in 100 µL of media were seeded in each well of a white-walled 96-well plate. After 24 h of incubation, cells were transfected with reaction mixtures containing 10 ng Nluc or Rluc reporter plasmid, 5 ng Fluc internal control reporter plasmid, and 85 ng effector protein expression plasmid. In some experiments where protein expression was titrated, the amounts of effector plasmid used are indicated in the figures and pcDNA3.1 HA plasmid was used to balance the total amount of plasmid transfected in each well to 100 ng. Transfection mixtures were prepared according to FuGENE HD protocol guidelines (Promega) and previously optimized conditions for luciferase reporter assays.^7,28^ Cells were harvested 48 h after transfection for protein isolation to confirm expression of effectors and reporter levels were measured by dual luciferase assay using a GloMax Discover luminometer (Promega).

Tethered function and altered specificity reporter assay data were analyzed as previously described.^7,11,28,63^ For each sample, a relative response ratio (RRR) was calculated by normalizing the Nluc or Rluc reporter light output, measured in relative light units (RLU), to the corresponding Fluc control output. The RRR values were then normalized to the respective negative control effector sample (HT-MS2-V5 in tethered function experiments and HT in altered specificity experiments). Repressive activity of each effector is presented as log2 Fold Change (log2 FC) in RRR relative to the negative control. For some experiments, as indicated in the figures, repressive activity of each effector was calculated as a percentage of WT effector activity. All reporter assays were performed in at least three independent experiments, each with three biological replicates. Data are presented as mean values +/- standard deviation overlaid with individual data points. Statistical analysis of data by two-tailed, unpaired Student’s t test or ordinary one-way ANOVA with post hoc tests for multiple comparisons was performed using GraphPad Prism 10 (GraphPad Software, Inc). Analysis methods and conventions for significance calling are noted in the figure legends and all data and statistics are reported in **Supplemental Table S2**.

#### Western blotting

Cells in wells of 96-well plates reserved for protein isolation were washed twice with 1x PBS and lysed in 20 µL radioimmunoprecipitation (RIPA) buffer (25 mM Tris-HCl pH 7.6, 150 mM NaCl, 1% NP-40, 1% sodium deoxycholate, 0.1% SDS) supplemented with 2x cOmplete Protease Inhibitor Cocktail (Roche). Triplicate wells were combined for each sample. Samples were homogenized and cleared by centrifugation at 21,000*g*, 10 min, 4 ℃. Protein concentrations for the resulting lysates were determined by detergent-compatible Lowry assay (BioRad). Equal masses or 10 µg of protein for each sample were then incubated with 5x loading dye at 100 ℃ for 10 min, separated by SDS-PAGE on a 4-20% Mini-PROTEAN TGX gel (BioRad), and transferred to Immobilon P membranes (Millipore). Blots were blocked for 1 h and incubated with primary antibody (as indicated in the figures and described below) for 1 h at room temperature or overnight at 4 ℃ with gentle rocking. Blots were then probed with HRP-conjugated secondary antibodies (described below) for 1 h at room temperature, incubated with enhanced chemiluminescent substrate (Millipore), and imaged using a chemiluminescence imaging system (Azure Biosystems).

#### Antibodies

All antibodies used for western blotting were diluted to the indicated concentrations in 1x PBS-T with 5% w/v nonfat dry milk according to the manufacturer’s instructions. Tethered function assay effector proteins were detected using mouse anti-V5 antibody (Invitrogen; catalog #: R960-25) diluted 1:5,000. Altered specificity assay effector proteins were detected using mouse anti-HT antibody (Promega; catalog #: G9211) diluted 1:2,000. In co-immunoprecipitation assays, endogenous CNOT1 was detected using rabbit anti-CNOT1 (CST; catalog #: 44613) diluted 1:1,000. For loading controls, blots were also probed with either rabbit anti-H3 (CST; catalog #: 4499) diluted 1:2,000 or rabbit anti-vinculin (Invitrogen; catalog #: 70062) diluted 1:1,000. Goat anti-mouse (Thermo Fisher Scientific; catalog #: 31430) diluted 1:5,000 or goat anti-rabbit (CST; catalog #: 7074) diluted 1,10,000 HRP-conjugated secondary antibodies were used to allow chemiluminescent detection of proteins.

#### In vitro pull-down assays

*In vitro* pull-down assays were performed to assess direct protein interactions between PUM1 RD3 regions of interest and the reconstituted, purified NOT module of the CCR4–NOT complex, as previously described.^28,30^ Briefly, StrepII and MBP-tagged PUM1 constructs were expressed in *E.coli* BL21 (DE3) Star cells (Thermo Fisher Scientific) and purified from the lysate using StrepTactin sepharose resin (IBA). Purified NOT module containing CNOT1 (aa 1833-2361), CNOT2 (aa 344-540), and CNOT3 (aa 607-753) ^40,64^ was then added to the resin and incubated for 1 h. Bound proteins were eluted and analyzed by SDS-PAGE and Coomassie blue staining.

#### Crosslinking and mass spectrometry

For crosslinking experiments, PUM1 RD3-RBD (589-1186) was purified from a construct in the pSMT3 vector with an N-terminal His6-SUMO tag. Briefly, the PUM1 protein was expressed in *E. coli* BL21(DE3) RILcodon+ cells (Agilent) at 16 °C overnight induced with 0.1 mM IPTG. The *E. coli* pellet was resuspended in a buffer containing 20 mM HEPES pH 8.1, 0.3 M NaCl, 20 mM imidazole, 5% v/v glycerol, 2 mM DTT and sonicated to lyse cells. The cleared cell lysate was incubated with 5 mL Ni-NTA resin (Qiagen) and eluted with buffer (20 mM HEPES pH 8.0, 0.3 M NaCl, 300 mM imidazole, 5% v/v glycerol, 2 mM DTT). The fusion protein was digested with Ulp1 protease at 6 μg/mL for 1 h at 4 °C. The PUM1 protein was further purified with a Hi-Trap Heparin column (Cytiva). Heparin column buffer A contained 50 mM HEPES pH 7.5, 0.5 mM TCEP, and buffer B contained an additional 1 M NaCl. The column was eluted with a gradient of increasing buffer B from 0% to 100%. Heparin column peak fractions were concentrated to 5 mL and loaded onto a HiLoad 16/60 Superdex 75 column (Cytiva) in a buffer containing 20 mM HEPES pH 8.1, 0.2 M NaCl, 1% v/v glycerol, 0.5 mM TCEP. NOT module containing CNOT1 (aa 1833-2361), CNOT2 (aa 350-540), and CNOT3 (aa 607-748) was purified as described previously.^59^ Purified proteins were flash frozen in liquid nitrogen in freezing buffer (25 mM HEPES pH 8.1, 0.2 M NaCl, 1% v/v glycerol, 0.5 mM TCEP). Aliquots of frozen protein were stored at -80 ℃ until use. Complex formation was performed by pre-incubating PUM1 RD3-RBD with 1 mM RNA (5’-AAAU<u>UGUACAUA</u>AGCC, with PRE underlined) at a molar ratio of 1:1.25 protein:RNA for 10 min, followed by the addition of NOT module at a molar ratio of 1:1 PUM:NOT for 1 h. Crosslinking of the PUM:NOT:RNA complex was then facilitated by the addition of bis(sulfosuccinimidyl)suberate (BS3) homobifunctional crosslinker (Thermo Fisher Scientific; catalog #: A39266) at a final concentration of 1 mM. After 30 min, the crosslinking reaction was quenched by the addition of 1 M Tris pH 7.5 to a final concentration of 50 mM for 15 min. Quenched reactions were submitted for mass spectrometry analysis.

Sample processing, mass spectrometry, and data analyses were performed essentially as described previously.^65^ Crosslinked mixture was digested by the addition of Trypsin/LysC Mix Mass Spec Grade (Promega) to a concentration of 0.033 μg/μL and incubation overnight at 40 °C. Digested samples were stored at -80 °C for subsequent mass spectrometry analysis.

Protein digests for the first sample replicate (MS1) were analyzed via LC/MS using a Q Exactive Plus mass spectrometer (ThermoFisher Scientific) interfaced with an M-Class nanoAcquity UPLC system (Waters Corporation) equipped with a 75 μm × 150 mm BEH C18 column (1.8 μm particle, Waters Corporation) and a C18 trapping column (180 μm × 20 mm) with 5 μm particle size. The second sample replicate (MS2) was analyzed using an Orbitrap Ascend mass spectrometer (ThermoFisher Scientific) coupled with a nanoVanquish UPLC system (ThermoFisher Scientific), which was equipped with an Easy-Spray™ PepMap™ Neo 2 μm C18 75 μm x 150 mm analytical column and a PepMap™ Neo 5 μm C18 300 μm x 5 mm trapping column. Five microliters of digested sample were injected onto the systems. The mass spectrometers were operated in positive ion mode with a spray voltage of 2000 volts. Both MS1 and MS2 analyses were conducted using the Orbitrap analyzer, and Higher-energy Collisional Dissociation (HCD) was used for fragmentation. Data and crosslinked peptides for MS1 and MS2 analyses are listed in **Supplemental Table S3**.

Intermolecular and intramolecular crosslinked residues were identified, and crosslinked residues were visualized by xiNET (crosslinkviewer.org).^66^ Modeling of crosslinked residues onto existing NOT module crystal structure was performed using Pymol (Schrödinger, LLC).

### Co-immunoprecipitation assays

Co-immunoprecipitation assays to assess the interaction between RD3 and the endogenous CNOT complex in cells were performed as previously described^28^ with minor protocol modifications. Briefly, 200,000 HCT116 cells in 2 mL of media were seeded in each well of a 6-well plate. 24 h post seeding, cells were transfected with reaction mixtures containing 3 µg effector protein expression plasmid. To achieve similar protein expression levels across effectors, 3 µg of plasmid for MS2 V5 and WT PUM1 RD3 and 1 µg for IDR muts was used, and pcDNA3.1 HA plasmid was added to balance the total amount of DNA transfected in each well to 3 µg as needed. Transfection mixtures were prepared according to FuGENE HD protocol guidelines (Promega) and previously optimized conditions for 6-well assays.^7,28^ 48 h after transfection, cells from four replicate wells for each effector were harvested into PBS, pooled together, and collected by centrifugation at 1,000*g*, 3 min, 4 ℃. Cells were then lysed and homogenized in 500 µL Buffer A (50 mM Tris-HCl pH 7.5, 500 mM NaCl, 0.5% Triton X-100, 1 mM EDTA) supplemented with 2x cOmplete Protease Inhibitor Cocktail (Roche). Cell debris was removed by centrifugation at 10,000*g*, 10 min, 4 ℃ and the resulting supernatant was then cleared through a 0.45-micron filter (Millipore) for 3 min at 4,000*g*. Lysate protein concentrations were determined by detergent-compatible Lowry assay (BioRad). For each sample, 500 µg of total protein in 400 µL of lysis buffer with 4 µg of RNase A (Promega) and 40 U of RNase ONE (Promega) was added to 8.3 µL of Protein A Dynabeads (Invitrogen) that were pre-incubated overnight with 2 µg of anti-V5 antibody. Samples were then incubated with end-over-end rotation at 4 ℃ for 2 h. Post-incubation, beads were washed six times with 500 µL of Buffer A and eluted in 5x SDS-PAGE loading dye. 40% of each co-IP sample was analyzed by western blot along with 10 µg of the input lysate. Successful RNase treatment was confirmed by isolating RNA from input and flow-through samples using Maxwell RSC simplyRNA Tissue Kit (Promega) with 2x DNase added and running equal volumes on a formaldehyde agarose gel.

## Data availability

All data are contained within the article.

## Supporting information

Supplemental Table S1 Plasmids and Sequences

Supplemental Table S2 Data and Statistics

Supplemental Table S3 Crosslinked peptides replicates MS1 and MS2

## Acknowledgements

We thank members of the Goldstrohm lab for helpful advice and comments including Dr. Robert Connacher and Dr. Katie McKenney. We thank Dr. Alex Holehouse and Dr. Kanishk Jain for insightful discussions on this research. The authors acknowledge use of the NIEHS Mass Spectrometry Research Center (ZIC ES103005). This research was supported by the National Institutes of Health grant R01 GM150468 to A.C.G; the Intramural Research Program of the National Institutes of Health, National Institute of Environmental Health Sciences ZIA-ES050165 to T.H. and ZIC ES103005 to J.W.; and the National Cancer Institute ZIA-BC011977 to E.V. E.B.D was supported by University of Minnesota’s Targets of Cancer Training Program, NIH grant T32 CA009138. This research was supported by the Intramural Research Program of the National Institutes of Health (NIH). The contributions of the NIH authors are considered Works of the United States Government. The findings and conclusions presented in this paper are those of the authors and do not necessarily reflect the views of the NIH or the U.S. Department of Health and Human Services.

## Author contributions

Elise B. Dunshee: investigation, formal analysis, methodology, conceptualization, resources, visualization, writing original draft, review, editing

Brenna Saladin: investigation, formal analysis, resources

David Turner: investigation, formal analysis, resources

Chen Qiu: investigation, formal analysis, resources, visualization

Robert Dutcher: investigation, formal analysis, resources, visualization

Jason Williams: investigation, formal analysis

Josh Corbo: resources, investigation

Liv Wolcott: resources

Amanda Korte: resources

Becky Haugen: resources

Traci Hall: conceptualization, funding acquisition, project administration, supervision, writing – review, editing

Eugene Valkov: conceptualization, funding acquisition, project administration, supervision writing – review, editing

Aaron C. Goldstrohm: conceptualization, funding acquisition, project leadership and administration, supervision writing original draft, review and editing

