## Supplemental Figures for "Human Pumilio proteins use fuzzy multivalent hydrophobic interactions to recruit the CCR4–NOT deadenylase complex to repress mRNAs"

### Supplementary Figure Legends

**Figure S1. PUM1 701-827 is sufficient for repression.** (A) Diagram of PUM1 RD3 and PUM1 RD3 701-827 effector proteins. Tethered function reporter assay is shown comparing repressive activity of RD3 to the truncated effector. N=9 (three experimental repeats, each with three biological replicates). All data are shown as mean and individual points +/- standard deviation of $\log_2$ Fold Change ($\log_2$ FC) values relative to negative control HaloTag (HT). Significance indicated above the x-axis denotes comparisons to HT, while significance indicated below the x-axis denotes comparisons between specific effectors. For significance calling, ns = not significant where $p \geq 0.05$, $p < 0.05$ = *, $p < 0.01$ = **, $p < 0.001$ = ***, $p < 0.0001$ = **** based on ordinary one-way ANOVA and Tukey's test for multiple comparisons. Western blot confirming expression of V5-tagged effector proteins is shown below. Vinculin served as a loading control.

**Figure S1**
**Dunshee et al**

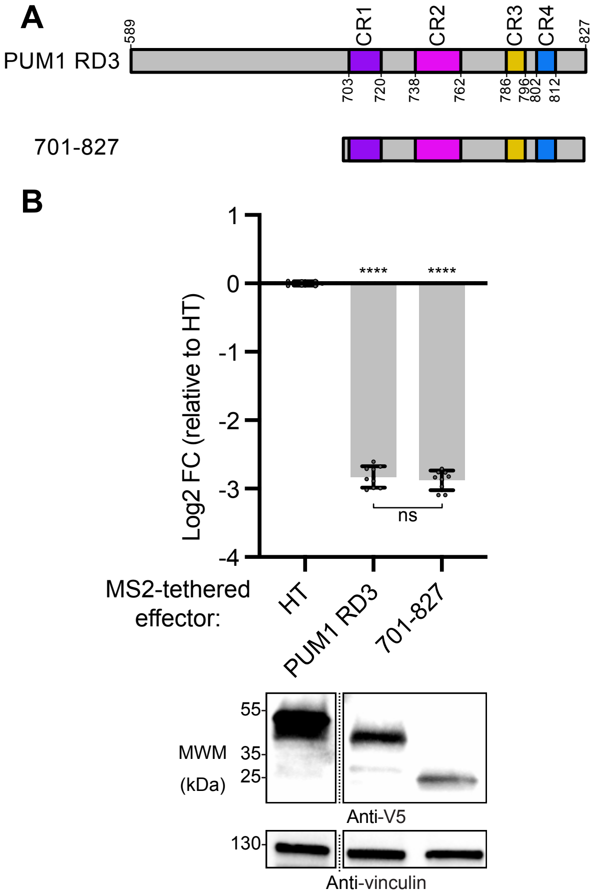

**Figure S2. Conserved regions of PUM2 RD3 are not necessary for repressive activity; Multiple regions of RD3 are functionally redundant.** (A) Tethered function reporter assay comparing repressive activity of WT PUM2 RD3 to RD3 effectors with individual and combined CR deletions. An RD3 effector with F683A and F690A substitutions was also tested. N≥9 (At least three experimental repeats, each with three biological replicates). All data are shown as mean and individual points +/- standard deviation of % Repressive Activity values relative to WT PUM2 RD3. Significance indicated denotes comparisons of each effector to WT. For significance calling, ns = not significant where $p \geq 0.05$, $p < 0.05$ = *, $p < 0.01$ = **, $p < 0.001$ = ***, $p < 0.0001$ = **** based on ordinary one-way ANOVA and Dunnet's test for multiple comparisons. Western blot confirming expression of V5-tagged effector proteins is shown below. Vinculin served as a loading control. (B) Tethered function reporter assay comparing the repressive activity of WT RD3 to truncated RD3 effectors and quarter deletions. N≥9 (At least three experimental repeats, each with three biological replicates). All data are shown as mean and individual points +/- standard deviation of % Repressive Activity values relative to WT PUM2 RD3. Western blot confirming expression of V5-tagged effector proteins is shown below. Vinculin served as a loading control.

**Figure S2**
**Dunshee et al**

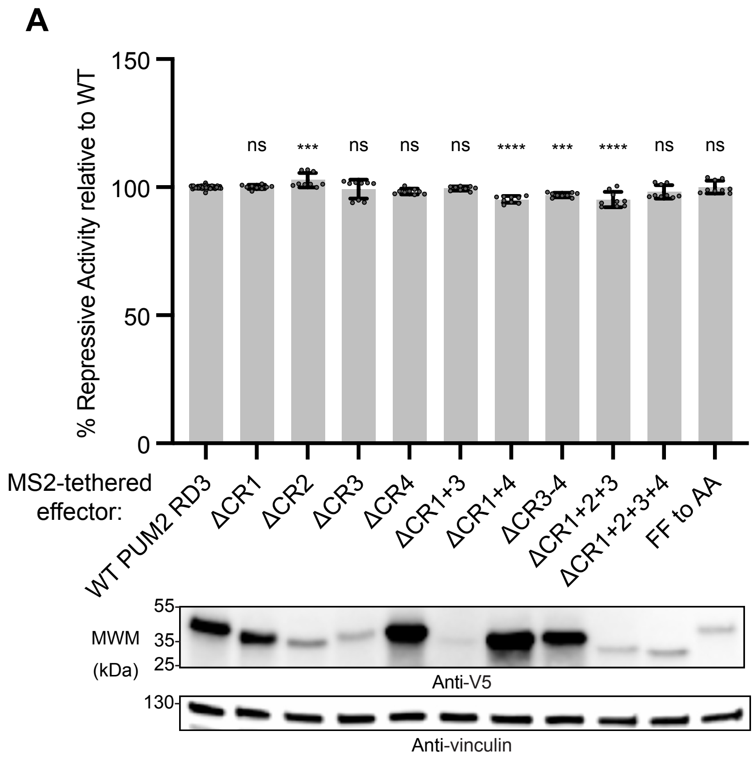


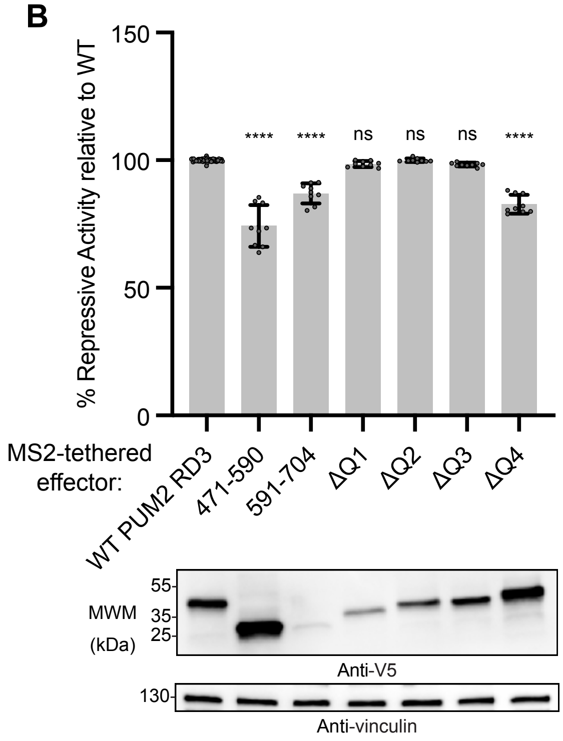

**Figure S3. PUM1 RD3-RBD:NOT module complex formation and crosslinking.** (A) Size exclusion chromatography UV traces of the ribonucleoprotein (RNP) complex formed with recombinant, purified PUM1 RD3-RBD and NOT module proteins along with PUM target RNA. UV absorbance measured in mAU at 254nm and 280nm is plotted on the y-axis. Peak fractionation intervals and corresponding volumes are shown on the x-axis. Traces for PUM1 RBD incubated with NOT module proteins and PUM target RNA are shown below. (B) SDS-PAGE and Coomassie blue staining of proteins in each collected fraction. (C) Analysis of BS3-crosslinked (XL) PUM1 RD3-RBD:NOT module complexes. SDS-PAGE and Coomassie blue staining of PUM1 RD3-RBD and NOT module proteins alone, in complex without crosslinking before and after clarification by centrifugation, and in complex crosslinked with BS3.

**Figure S3**
**Dunshee et al**

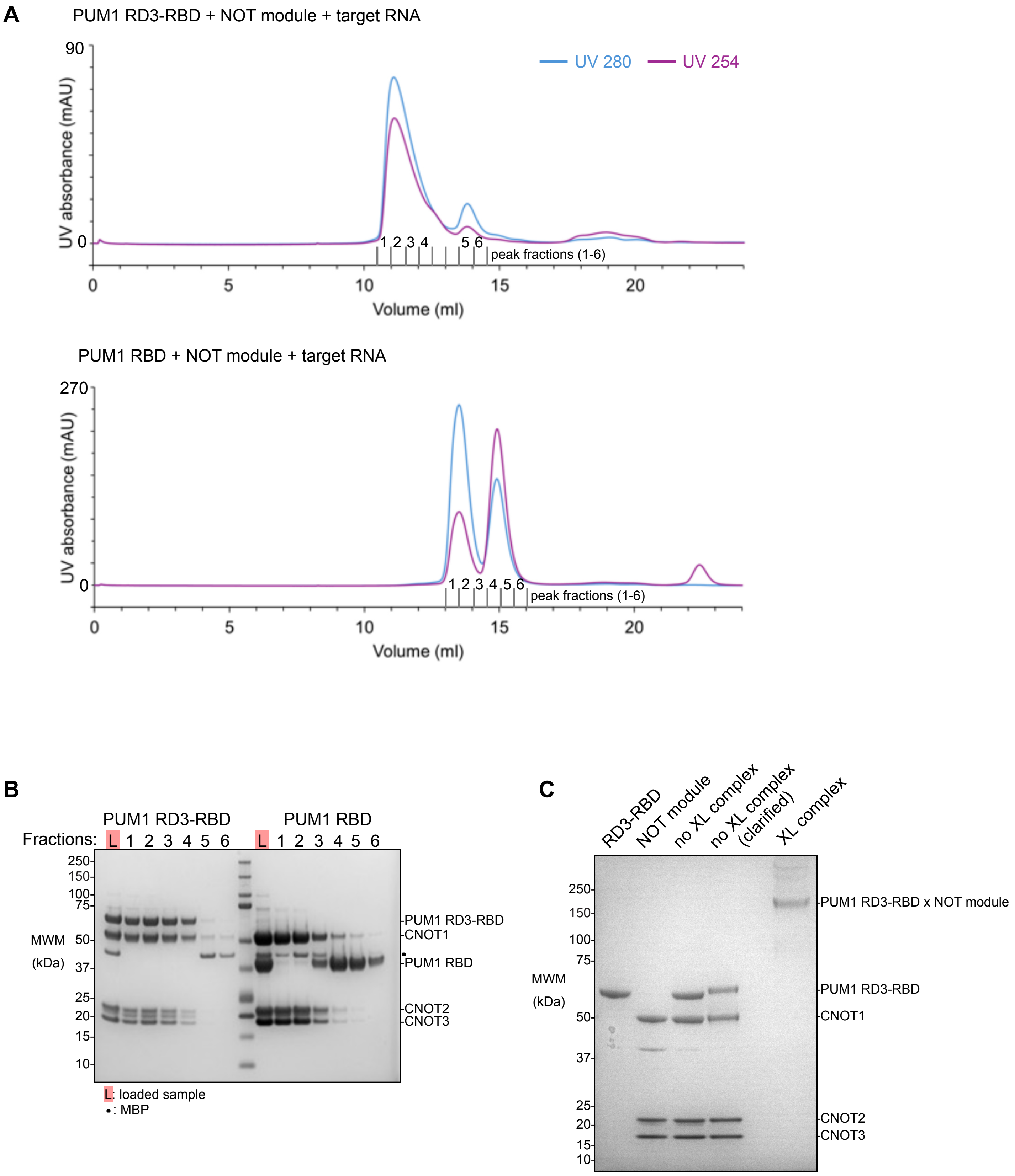

**Figure S4. The activity of RD3 repressive peptides withstands targeted deletions or substitutions.** Tethered function reporter assay comparing repressive activity of PUM1 RD3 minimal repressive peptides to RD3 effectors with deletions or substitutions made within these regions. CR deletions and FF to AA substitutions in the context of WT RD3 are described in **Figure 3.** The P789A substitution was tested based on its potential to disrupt a structural modeling-predicted interaction interface, predicted using AlphaFold, between the RD3 774-800 region and the NOT module. N=9 (three experimental repeats, each with three biological replicates). All data are shown as mean and individual points +/- standard deviation of $log_2$ Fold Change ($log_2$ FC) values relative to negative control HaloTag (HT). Significance indicated above the x-axis denotes comparisons to HT, while significance indicated below the x-axis denotes comparisons between specific effectors. For significance calling, ns = not significant where $p \geq 0.05$, $p < 0.05$ = *, $p < 0.01$ = **, $p < 0.001$ = ***, $p < 0.0001$ = **** based on ordinary one-way ANOVA and Tukey's test for multiple comparisons. Western blot confirming expression of V5-tagged effector proteins is shown below. Vinculin served as a loading control.

**Figure S4**
**Dunshee et al**

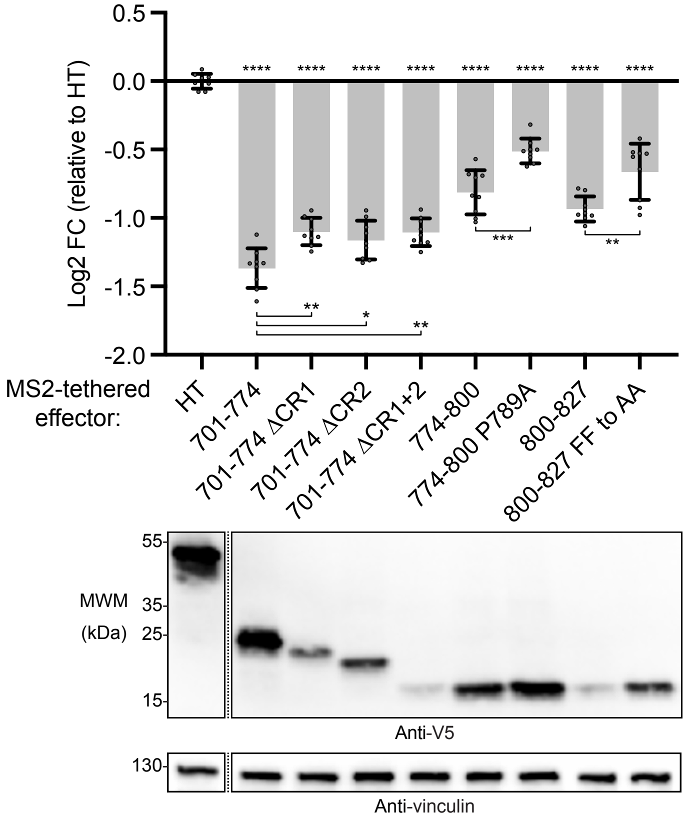
